# Satellite microglia-like cells in human dorsal root ganglia and changes with diabetic neuropathy

**DOI:** 10.64898/2026.05.12.724479

**Authors:** Khadijah Mazhar, Jayden A. O’Brien, Michael A. Wilde, Harikrushnaa Srikanth, Andi Wangzhou, Victoria Pastor, Charlynn W. Maina, Nora S. Arefin, Marisol Mancilla Moreno, Ishwarya Sankanarayanan, Diana Tavares-Ferreira, Theodore J. Price

## Abstract

Phagocytic and immune-like cells have been observed in the satellite envelope of neuronal somata in peripheral sensory ganglia of many species for several decades. These cells likely play an important role in normal function of sensory neurons and they may also play an important role in neuronal dysfunction and neurodegeneration seen with neuropathy. Recent findings have described a satellite macrophage population transcriptomically similar to microglia in peripheral ganglia of some mammalian species. The function of these cells, and the mechanisms by which they may influence neurons in neuropathy are unclear. We sought to understand the phenotype and localization of these cells in the human dorsal root ganglion (hDRG) using large-scale single nucleus and spatial transcriptomic datasets from individuals with and without a history of peripheral diabetic neuropathy. We observed a large population of macrophages that express classical microglia makers such as *TMEM119* and *P2RY12* in the hDRG, as previously described. Our findings confirm that these microglia-like cells (MLCs) localize to the satellite envelope around neuronal somata, yet are transcriptomically distinct from all glial cell types characterized in the hDRG. These MLCs exhibit changes in abundance and localization with diabetic painful neuropathy (DPN) in both the hDRG and sural nerves suggesting that they are not exclusively localized to the DRG. We conclude that microglia-like cells are likely the resident tissue macrophage (RTM) of the hDRG, and perhaps the peripheral nervous system (PNS) given their localization to the sural nerve and other ganglia, where they are predicted to regulate homeostatic neuronal functions and response to injury.

**Highlights:**

- MLCs are likely the RTM of hDRGs
- MLCs localize to the satellite envelope and recede with Nageotte nodule formation
- MLC activation state and signaling shift with diabetic neuropathy
- MLCs are also present in other ganglia and sural nerve

## Introduction

Satellite glial cells (SGCs) are specialized glial cells in the peripheral nervous system that closely encase dorsal root ganglion (DRG) neurons. These cells are recognized to play an important role in pain and other sensory aspects of peripheral neuropathy in rodent models; however, several recent studies of human DRG (hDRG) have suggested that SGCs are intertwined with a macrophage or microglial-like cell type^1^ that may actually reside in closer to proximity to somatosensory neurons than SGCs^2^. In the most detailed of these recent works, Wu et al., describe a resident macrophage population in large animals transcriptomically similar to central nervous system (CNS) microglia in peripheral sensory and sympathetic ganglia^2^. Single-cell RNA-sequencing analysis shows that this population – termed peripheral nervous system (PNS) microglia-like cells (MLCs) – clusters with CNS microglia rather than marrow-derived macrophages (MDMs), expresses microglia markers such as *P2RY12* and *TMEM119*, and does not express some MDM genes such as *MRC1* and *LYVE1*. Single-cell transcriptomics and immunostaining of human embryonic tissues from yolk sac, CNS, and DRG trace the emergence of microglia and MLCs, and demonstrate that they share a common macrophage progenitor that migrates to the nervous system from the yolk sac. Wu et al. also show that the MLCs primarily reside within the SGC envelope around neuronal somata and suggest that MLCs may regulate glial and neuronal function and phenotype. Hunt et al. also report macrophages expressing microglia marker *CX3CR1* that form interconnected networks around neuronal somata in the satellite envelope^1^.

Although the notion of resident microglia-like cells in the PNS appears to be a new concept, a detailed review of the literature reveals that these cells have been detected previously using other methods that predate single cell and spatial RNA sequencing. The first mention of ‘microglia-like cells’ comes from Fiori et al. in 1981 who describe satellite cells with electron-dense, elongated cell bodies that lack basal lamina in chick ciliary ganglia^3^. They observe that these cells resemble CNS microglia morphologically and are distinct from satellite glial cells. Since the work of Fiori and colleagues, several others have also observed immune marker-expressing and phagocytic cells in the satellite envelope of neuronal somata in peripheral ganglia (Table 1). These studies include imaging of peripheral ganglia using light, confocal, transmission electron, and scanning electron microscopy, as well as staining for different macrophage and microglial markers. They encompass a variety of species and demonstrate that, for some species such as rodents, these cells are detected only after injury. Fascinatingly, authors have come to different conclusions regarding the identities of satellite envelope cells with immune phenotypes, such as whether they are macrophages or glia, yet the original data and qualitative observations reported in these papers are largely consistent^1–24^.

**Table 1.** Previous work demonstrating satellite macrophages in peripheral ganglia. Green rows indicate studies that suggested the presence of macrophages similar to CNS microglia. DRG = dorsal root ganglia. SCG = superior cervical ganglia. TG = trigeminal ganglia. EM= electron microscopy. LM = light microscopy. IHC=immunohistochemistry. SGCs = satellite glial cells.

| Author | Year | PMID | Ganglia | Species | Observations |
| --- | --- | --- | --- | --- | --- |
| Matthews | 1972 | 4402334 | SCG | Rat | EM: phagocytic sheath cells around neurons after injury |
| Smith | 1972 | 5011947 | DRG | Mouse | LM: mononuclear leukocytes among satellite cells after injury |
| Hill | 1974 | n/a | DRG | Rat | Scanning EM: macrophages around somas are visually distinct from SGCs |
| Fiori | 1981 | 7242910 | Ciliary | Chickens | LM/EM: Electron-dense basal lamina(-) **"Microglia-like cells"** that are morphologically similar to CNS microglia and distinct from electron-lucent basal lamina(+) SGCs in the satellite sheath |
| Esiri | 1989 | 2754017 | Multiple | Human | IHC: satellite cells express MCH II and some express macrophage markers. Observed TG, DRG, SCG, Celiac ganglia, and sympathetic chain ganglia |
| Scaravilli | 1991 | 1919607 | DRG | Human (0-80 yrs), Rat | IHC: resident satellite macrophages present in humans and not rats<br>EM (humans): satellite cells include electron-lucent basal lamina(+) SGCs and electron-dense basal lamina(-) cells. Nageotte nodule cells appear to be SGCs or Schwann cells<br>EM (rat): SGCs are flat and the envelope is only one layer. No Nageottes, but different types of cells in layers around degenerating neurons |
| Glenn | 1993 | 7507913 | TG | Rat | Lectin histochemistry: ramified macrophages resembling microglia and positive for microglial stains are seen throughout the ganglia |
| Lu | 1993 | 8315414 | DRG | Rat | LM/IHC: Satellite macrophages appear after injury that look like SGCs but are S-100 negative |
| Schreiber | 1995 | 7658197 | SCG | Rat | IHC/EM: Satellite macrophages seen after injury |
| Hu | 2003 | 14769352 | DRG | Rat | IHC: Macrophages seen beneath capsule around somas after injury |
| Dubovy | 2007 | 17931774 | DRG | Rat | IHC: Satellite macrophages appear after injury in both ipsi- and contra- DRGs |
| Patro N | 2010 | 20455319 | DRG | Rat | IHC: satellite 'microglia' that change morphologically after injury like microglia |
| Niemi | 2013 | 24107955 | SCG | Rat | IHC/PCR/Gene knockout: Macrophages appear around neuronal somas after injury and promote neurite growth after Wallerian degeneration |
| Ton | 2013 | 23701965 | DRG | Rat | IHC: Resident macrophages ring around somas after nerve ligation but less so with STZ/diabetic model. |
| Vega-Avelaira | 2013 | 20003309 | DRG | Rat | Microarray/IHC: Resident macrophages accumulate around large diameter somas after injury in adult but not young rats |
| Koeppen | 2016 | 27142428 | DRG | Human | IHC: Satellite monocytes that may penetrate neuronal membrane, and may be in residual nodules in Friedreich ataxia |
| Avraham | 2021 | 34586065 | DRG | Mouse | scRNA-seq: Some macrophages express the microglia markers Tmem119, P2ry12, and Trem2 |
| Iwai | 2021 | 34645458 | TG | Mouse | IHC/transgenic mice: resident macrophages contacted neurons after injury |
| Feng | 2023 | 36763532 | DRG | Mouse | scRNA-seq/fate-mapping: There are Tmem119+ microglia-like cells and Iba1+ SGCs after injury |
| Guimarães | 2023 | 37254842 | DRG | Mouse | Proliferation of resident CX3CR1+ macrophages that closely associate with somas after injury promote neuropathic pain |
| Grosu | 2024 | 39581686 | DRG | Mouse | After SNI, CX3CR1+ macrophages form perineuronal rings around CSF1+ DRG neurons; incidence varies across L3–L5 and neuronal subtypes. |
| Sapio | 2024 | 38691673 | DRG | Human | IHC/RNAscope: Shows satellite TMEM119+P2RY12+ADORA3+ and FABP7-macrophages |
| Shiers | 2025 | 40325011 | DRG | Human | IHC/RNAscope/VISIUM: CD68+ cells are seen around neurons as well as in Nageotte nodules in DPN DRGs. Some of the CD68+ cells are also FABP7+ |
| Wu | 2025 | 40199320 | DRG, SCG | Human, many | snRNA-seq/ATAC-seq/IHC: Microglia-like cells that share CNS microglia progenitor reside within/beneath SGC envelope of larger animals and not smaller ones |
| Hunt | 2026 | n/a | DRG | Human | IHC: CX3CR1+ macrophages seen associating with SGCs around neuronal somata and CD163+ macrophages were spread throughout the DRG |

Given the findings of several recent studies on DRGs from multiple species, it is likely that there are multiple cell types in the satellite envelope that express immune cell genes and perform immune functions in the hDRG. Some of these studies indeed report a diversity of cell types in the satellite envelope, and antigen presentation markers are present on both cells that do and do not express macrophage markers^25^. Even among cells that express macrophage markers, there is some confusion on whether they are of glial or macrophage lineage, as there are multiple reports of SGCs expressing macrophage genes^20,26,27^. Here, we aim to assess the possibility that MLCs are the primary immune cell and phagocyte of the satellite envelope in hDRG. We incorporated published single-nuclei and spatial transcriptomic data from hDRG^7,28–31^ and other PNS tissue^32–37^ to support this hypothesis, showed that MLCs are likely not of glial or marrow-derived macrophage lineage, and evaluated changes in MLC activation state and signaling with DPN. We conclude that these MLCs are likely resident macrophages of the hDRG, and potentially of the PNS more broadly.

## Results

### Human DRG single-nuclei data reveals a population of macrophages expressing microglial markers

Published single-nuclei transcriptomic data of human dorsal root ganglia ^28–31^ from 87 donors aged 2 months - 88 years old and spanning cervical, thoracic, and lumbar spinal levels (Table S1) provided 450,570 cells for analysis (Figure 1A and S1A). A cluster of cells enriched in *CSF1R*, a marker for the family of mononuclear phagocytes (MNPs), was identified and further subclustered (Figure 1B). The 11 clusters identified included six groups of likely monocyte-derived MNPs: 1) dendritic cells that were *CLEC10A*+ *CD1D*+ (Figure 1C); 2) monocytes (*CD300E*+); 3) likely differentiating monocytes that expressed monocyte markers (*LYZ* and *S100A9*) and MDM markers (*MRC1* and *SIGLEC1*), and were enriched in differentiating monocyte genes *C1QA* and *MARCO*; 4) a late monocyte or early MDM group that was *LYZ+ S100A9+ MARCO+ MRC1+ SIGLEC1+* and enriched in *GPNMB*; 5) a likely early MDM cluster enriched in *CD209* and *LILRB5*; and 6) a *MRC1+ SIGLEC1+* cluster of mature MDMs with very low expression of monocyte and immature macrophage markers.

**Figure 1.**
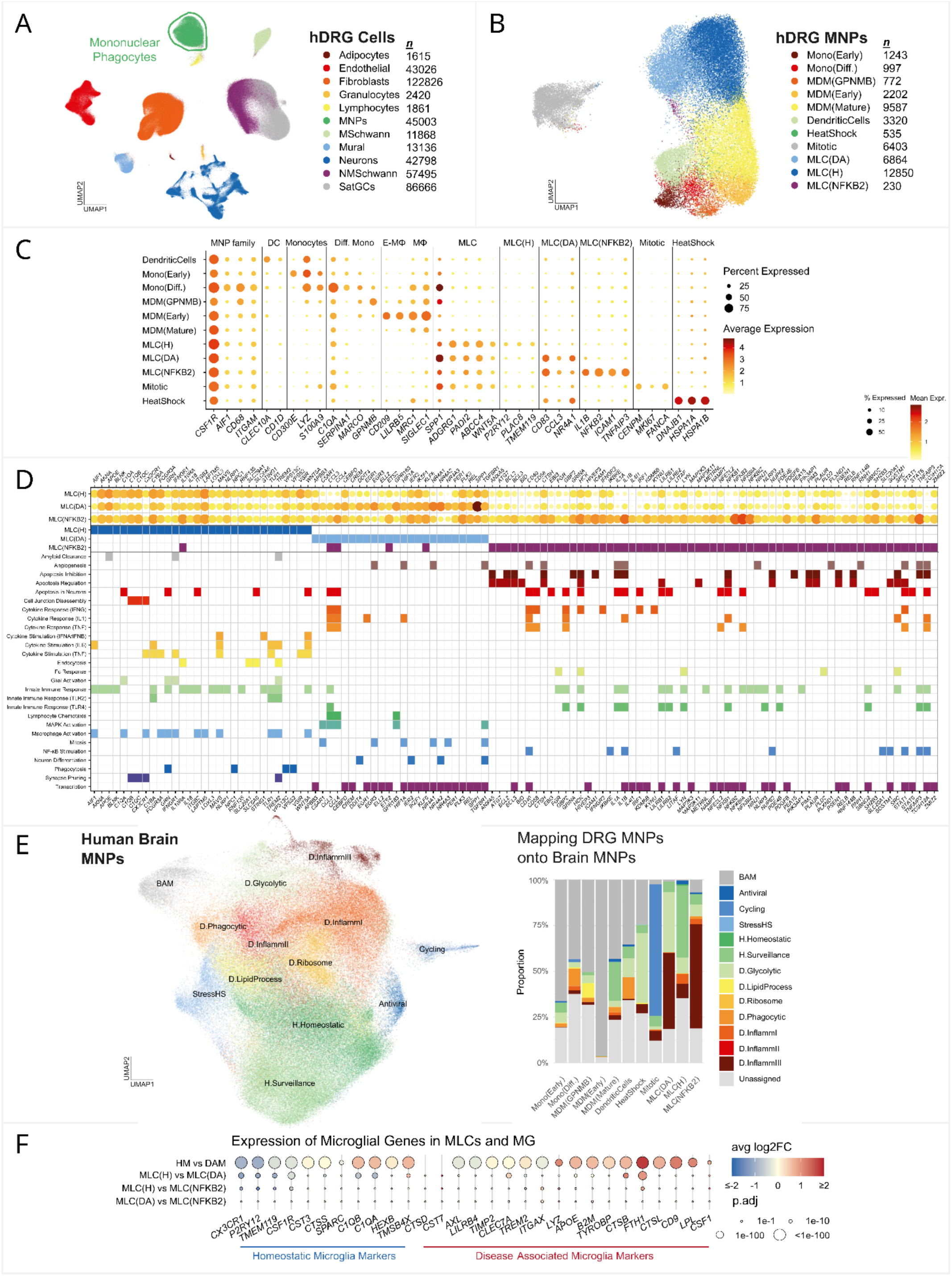
Identification of MLCs in human DRG transcriptomic data. A) UMAP of human DRG cells showing a cluster of MNPs and B) MNP subclusters. C) Dot plot of gene expression in MNP cell types shows that MLCs and monocyte-derived macrophages had distinct markers. D) Functional categories associated with genes enriched in each MLC type (padj < 0.05). E) Mapping DRG MNPs onto a dataset of human brain MNPs (left) predicted which CNS macrophages DRG MNPs are most like. Predictions are shown as % of cells that mapped to different brain MNPs per DRG MNP cell type (middle, right). F) Relative enrichment of known homeostatic and DAM markers in brain microglia (top row) and DRG MLC populations; regulation direction represents expression in the first group compared to the second group.

There were three clusters of MLCs enriched for the microglial gene *ADGRG1*^38^: 1) the largest, MLC(H), was enriched in homeostatic microglia genes *P2RY12*, *TMEM119,* and *CX3CR1* (Figure 1C); 2) another cluster, MLC(DA), was enriched in disease-associates microglia genes *CD83, CCL3,* and *SPP1*; 3) a small cluster of MLCs enriched in *NFKB2* and genes NFKB2 is known to induce (such as *IL1B*) was also observed close to the MLC(DA) cluster. As this MLC(NFKB2) group also expressed *CD83* and *CCL3*, it may reflect MLC(DA) cells in a further activated state and MLC(DA), in turn, may reflect an altered state compared to MLC(H) rather than a separate cell type. All groups of MLCs clustered separately from the likely marrow-derived *MRC1+* macrophages, consistent with Wu et al.^2^ While the MLC(H) group matches all of the transcriptomic characteristics by which Wu et al. describe microglia-like cells, the MLC(DA) may not have been identified as a separate group because they are transcriptomically similar to MLC(H) and are likely combined into one cluster in smaller datasets. Additionally, we found that MLCs were enriched in neurotrophic factor *WNT5A*, myelin citrullination enzyme *PADI2*, and prostaglandin transporter *ABCC4* relative to MDMs.

The last two MNP clusters were 1) a group of mitotic cells enriched in *MKI67* that clustered away from the other MNPs and 2) cells that expressed heat shock response genes such as *HSPA1A* that were scattered on the UMAP among the other MNP cell types (Figure S1A). As the mitotic cluster had greater expression of MLC markers *ADGRG1* and *WNT5A* compared to MDM markers *MRC1* and *SIGLEC1* (Figure 1C), we assessed whether MLCs are overrepresented among mitotic cells and found that while MLCs were 52% of non-mitotic MNPs, they were 81% of mitotic MNPs (Figure S1B). These observations suggest MLCs are a major class of MNPs in the hDRG and are more likely to proliferate than MDMs.

### MLC subpopulations align with homeostatic and disease-associated microglia

To identify potential functional differences between the three MLC clusters, enriched genes per cluster (padj. < 0.05) were intersected with Gene Ontology Biological Process terms^39^ through Enrichr^40^. The top 30 terms per MLC type were collectively sorted into 30 high-level categories (Table S2). 27 selected categories and their associated genes (Figure 1D) predicted that the MLC subpopulations may play slightly different roles with respect to homeostatic functions, cytokine signaling, and innate immune response. MLC(H) expressed higher levels of genes involved in amyloid clearance, synapse pruning, glial activation, endocytosis, phagocytosis, and stimulation of type I interferon, IL6, and TNF production. MLC(DA) also expressed these genes and showed further enrichment of genes involved in lymphocyte chemotaxis, MAPK activation, mitosis, and neuron differentiation. MLC(NFKB2) was enriched in genes that may regulate apoptosis, including inhibiting apoptosis, and that respond to activation of Fc receptors during opsonization. Both MLC(DA) and MLC(NFKB2) were enriched in genes that promote transcription, angiogenesis, and response to type II interferon, TNF, and IL1. MLC(H) and MLC(NFKB2) were predicted to be enriched in innate immune response genes with MLC(NFKB2) indicating TLR4 activation. As MLC(H) enriched genes were largely not specific, MLC(DA) exhibited greater enrichment in genes compared to MLC(H), and MLC(NFKB2) had the highest enrichment, these groups may reflect different activation states. MLC(DA) may reflect activation of additional transcriptional programs relative to MLC(H) and MLC(NFKB2) may reflect the highest level of activation in response to inflammation.

To explore the idea that the different subclusters of MLCs may be analogous to major groups of microglia in the CNS, hDRG MNPs were compared with human brain MNPs using a dataset by Sun et al.^41^ that included CNS microglia and brain-associated macrophages (BAMs) (Figure 1E and S1C). Of the 12 microglial states Sun et al. describe, we found that the Neuronal Surveillance and Homeostatic groups were relatively enriched in homeostatic microglia (HM) genes *CX3XR1*, *P2RY12*, and *TMEM119* while the other 10 groups were relatively enriched in disease-associated microglia (DAM) genes *CTSB* and *FTH1,* with the Glycolytic and InflammatoryIII groups showing highest enrichment (Figure S1E). Mapping of hDRG MNPs onto brain MNPs showed that hDRG MDMs were most like marrow-derived BAMs, and that MLCs were most similar to the microglia. MLC(H) were more similar to the Neuronal Surveillance HM than any other microglia type. MLC(DA) and MLC(NFKB2) were more similar to the InflammatoryIII DAM than any other microglia type, with a significant proportion of MLC(DA) also mapping to the Glycolytic DAM. Many hDRG MNPs were unassigned as they did not map confidently to any CNS MNP (prediction score < 0.5), suggesting that there are also major differences between hDRG MNPs and their CNS correlates.

We also evaluated MLC enrichment of genes that can help identify microglial state in the HM-DAM transition in mice^42^ (Figure 1F). MLC(H) were enriched in the key HM genes *CX3XR1*, *P2RY12*, and *TMEM119* along with some other HM genes such as *CSF1R*. MLC(DA) and MLC(NFKB2) were enriched in DAM genes *CTSB* and *FTH1* relative to MLC(H), and there were few differences between MLC(DA) and MLC(NFKB2) suggesting that both groups may represent disease-associated states of MLCs. Interestingly, MLC(H) exhibited more differences with MLC(DA) than MLC(NFKB2), suggesting MLC transition may not be a linear process or that MLC(NFKB2) may not be the most activated or disease-associated state. If MLC(NFKB2) is the most activated state, it could be that its transition from MLC(DA) may restore some phenotypes of MLC(H) that were lost with transition of MLC(H) to MLC(DA). Additionally, while HM genes *C1QA/B* were enriched in MLC(H) compared to MLC(DA) in hDRG, they were enriched in DAMs compared to HM in the brain, suggesting there may be some differences in the MLC(H)-MLC(DA) transition compared to the HM-DAM transition. There are likely species differences in this transition as well, as mouse HM marker *TMSB4X* was enriched in both human DAMs and hDRG MLC(DA) and several mouse DAM markers did not show enrichment in either DAMs or MLC(DA) compared to their homeostatic counterparts. Olah et al. have also observed that DAM marker genes may be different in humans compared to mice^43^.

### MLCs are likely not marrow-derived and arise in the DRG during development

As MLCs cluster more closely with other macrophages than any other hDRG cell type, we used trajectory analysis to assess the likelihood that MLCs may differentiate from the other macrophages. The Slingshot algorithm^44^ was applied to the MNPs and determined that the MDMs and dendritic cells were transcriptomically linked to monocytes, supporting the notion that these cells are of marrow-derived lineage (Figure 2A). The MLCs were not linked to the MDM family, and there is no identifiable cluster of cells that could represent their precursor in this dataset. As MLCs are thought to be a resident tissue macrophage (RTM) class, whose progenitors migrate to target organs from embryonic tissues during development^2^, we analyzed a human embryonic DRG dataset by Lu et al.^45^ to evaluate the presence of MLC precursors in the developing DRG. We identified 10 MNP clusters, of which 2 expressed MLC marker *WNT5A* and were enriched in *CX3CR1* and *P2RY12* (Figure 2B). One of the MLC clusters expressed MDM markers *MRC1, SIGLEC1,* and *LILRB5,* consistent with Wu et al.’s description of *P2RY12+MRC1+* MLC precursors. We also observed that mature MLCs were not present in gestational week (GW) 7, were less prevalent than MLC precursors in GW 8, and were more prevalent than precursors in GW 9-21 (Table S2). Wu et al. had also shown that MLC precursors peak in the DRG prior to 8 weeks of development and can be detected at low levels up to post-conceptual week 24.

**Figure 2.**
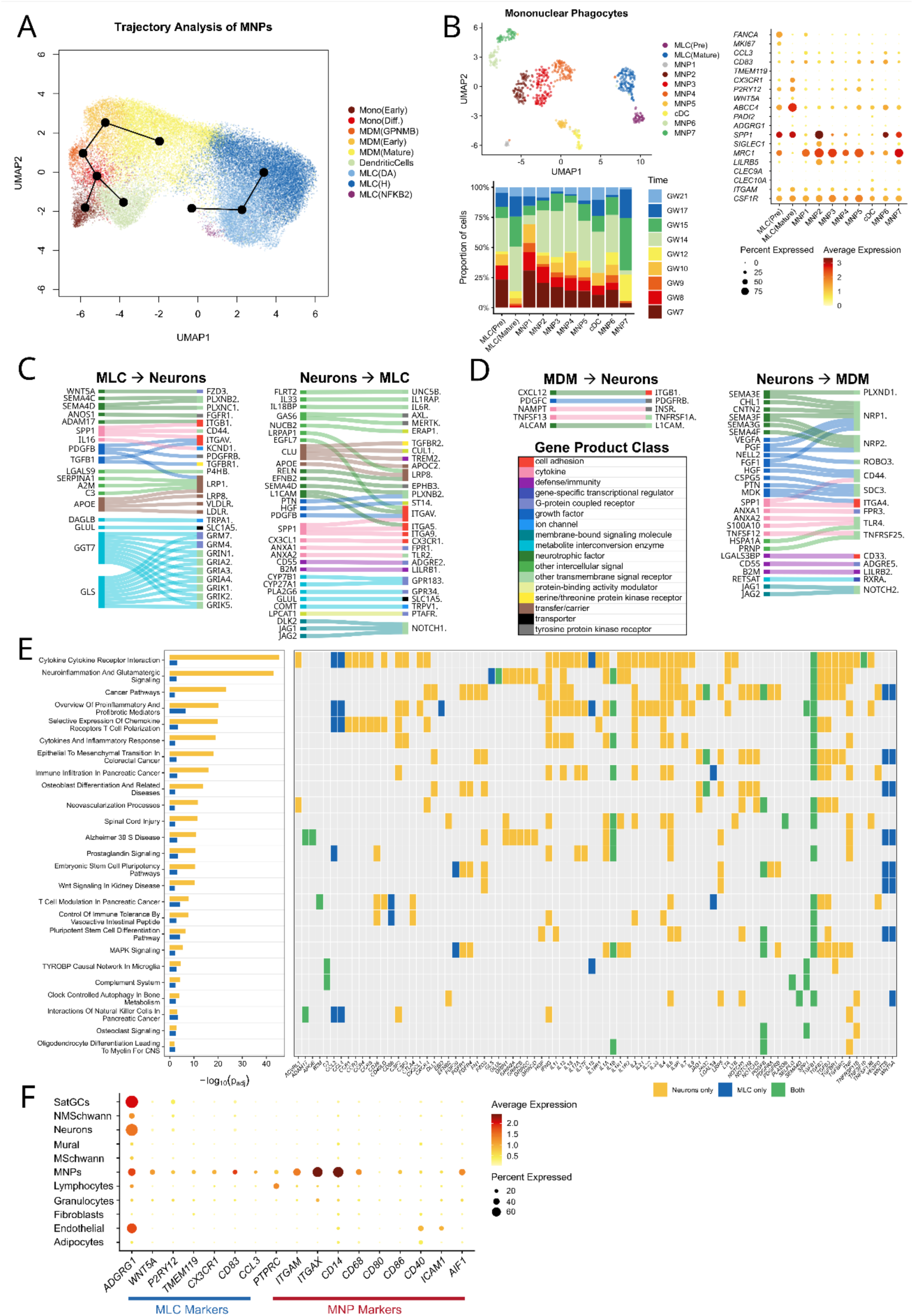
MLCs have lineage and functions distinct from MDMs and other DRG cell types. A) Trajectory analysis suggests that MDMs and dendritic cells are derived from monocytes while MLCs are not. B) *P2RY12+MRC1+* MLC precursors are found among human embryonic DRG MNPs. C) MLC-neuron interactome of ligand genes enriched in MLCs compared to MDMs and corresponding receptors expressed by DRG neurons (left) as well as receptor genes enriched in MLCs and corresponding ligands genes expressed in neurons (right). D) Interactome of signaling mechanisms predicted to be enriched in MDMs. For C and D, top interactions are shown (where normalized expression of neuronal genes > 0.1). E) Selected WikiPathways terms (p<0.01) enriched in the complete set of MLC signaling genes and the set of corresponding neuronal genes. F) Expression of MLC markers and MNP markers across all DRG high-level cell types to determine specificity of markers to MNPs and likelihood of glial expression.

### MLCs and MDMs are predicted to have differences in signaling with neurons

To assess how MLCs may differ functionally from MDMs, we predicted signaling mechanisms that may be enriched in either cell type. Genes that were differentially expressed between MLCs and MDMs (padj. <0.05) were intersected with a ligand-receptor database^46^ to identify specific signaling genes. Neuronal gene expression data was intersected with the database to identify ligand and receptor genes that pair with the signaling genes enriched in MLCs and MDMs. An interactome was generated of 308 ligand-receptor pairs between MLCs and neurons (Table S3), of which the top interactions (where neuronal gene normalized expression > 0.1) are shown (Figure 2C, Table S3). Similarly, an interactome of 142 signaling pairs was identified between MDMs and neurons (Figure 2D, Table S3).

MLCs may interact with neurons through diverse mechanisms. Interactome analysis predicted that cytokine signaling and activation of complement receptors may promote neuronal sensitization. It also suggested that MLCs may release small molecules that can directly stimulate neurons such as glutamate (represented in the interactome with glutaminase (*GLS*)) and arachidonic acid metabolites (represented by diacylglycerol lipase (*DAGLB*)). They could also protect neurons from overstimulation by converting glutamate to glutamine with glutamine synthase (*GLUL*) or provide glutamine for uptake by neurons with the transporter encoded by *SLC1A5.* MLCs may regulate neuronal or nerve growth through growth and neurotrophic factors, such as WNT5A, Anosmin-1 (*ANOS1*), TGFB1, and PDGFB. They may also stimulate autophagy and lysosomal activity with APOE and LRP1 related signaling.

When compared with signaling to MDMs, neuronal signaling to MLCs includes several interactions involving cell adhesion with integrins. Neurons may suppress immune functions by activating leukocyte immunoglobulin-like receptors (LILRBs) in both classes of macrophages. LILRB1 activation in MLCs may prevent phagocytosis^47^ and LILRB2 activation in MDMs may inhibit effector T cell activation^48^. Neurons could also promote phagocytosis mechanisms in MLCs such as efferocytosis with activation of AXL^49^ and ADGRE2^50^, amyloid clearance with TREM2^51^, and clearance of complement-bound particles with CD11 molecules (*ITGAM, ITGAX, ITGB1*). They may also stimulate MLC proliferation through CSF and CCN growth factors, while TLR2 activation could stimulate cytokine production-such as IL6 and TNF^52^-followed by cell death^53^. Neurons are predicted to signal to NRP1 on MDMs with SEMA3 molecules, which may be anti-inflammatory cues^54–57^, while IFN-γ signaling to MLCs may suppress NRP1 expression^56^. In contrast, neuronal signaling with SEMA4D to PLXNB2 on MLCs may be pro-inflammatory^58,59^. MLCs may also signal with SEMA4D, which may inhibit neuronal growth and promote neurodegeneration^60^.

Interactome genes enriched in MLCs compared to MDMs were analyzed with the WikiPathways^61^ library in Enrichr^40^ - including both ligand and receptor genes that are enriched relative to MDMs – and predicted that these signaling genes may promote adhesion with neurons, regulate neuronal development and apoptosis, promote neuroexcitation and inflammation, and receive regulation for phagocytosis (Figure 3E, Table S2). All of the predicted pathways were recapitulated when performing gene set enrichment analysis on the neuronal genes in the MLC-neuron interactome (Figure 3E, Table S2), supporting these predicted functions for MLCs.

**Figure 3.**
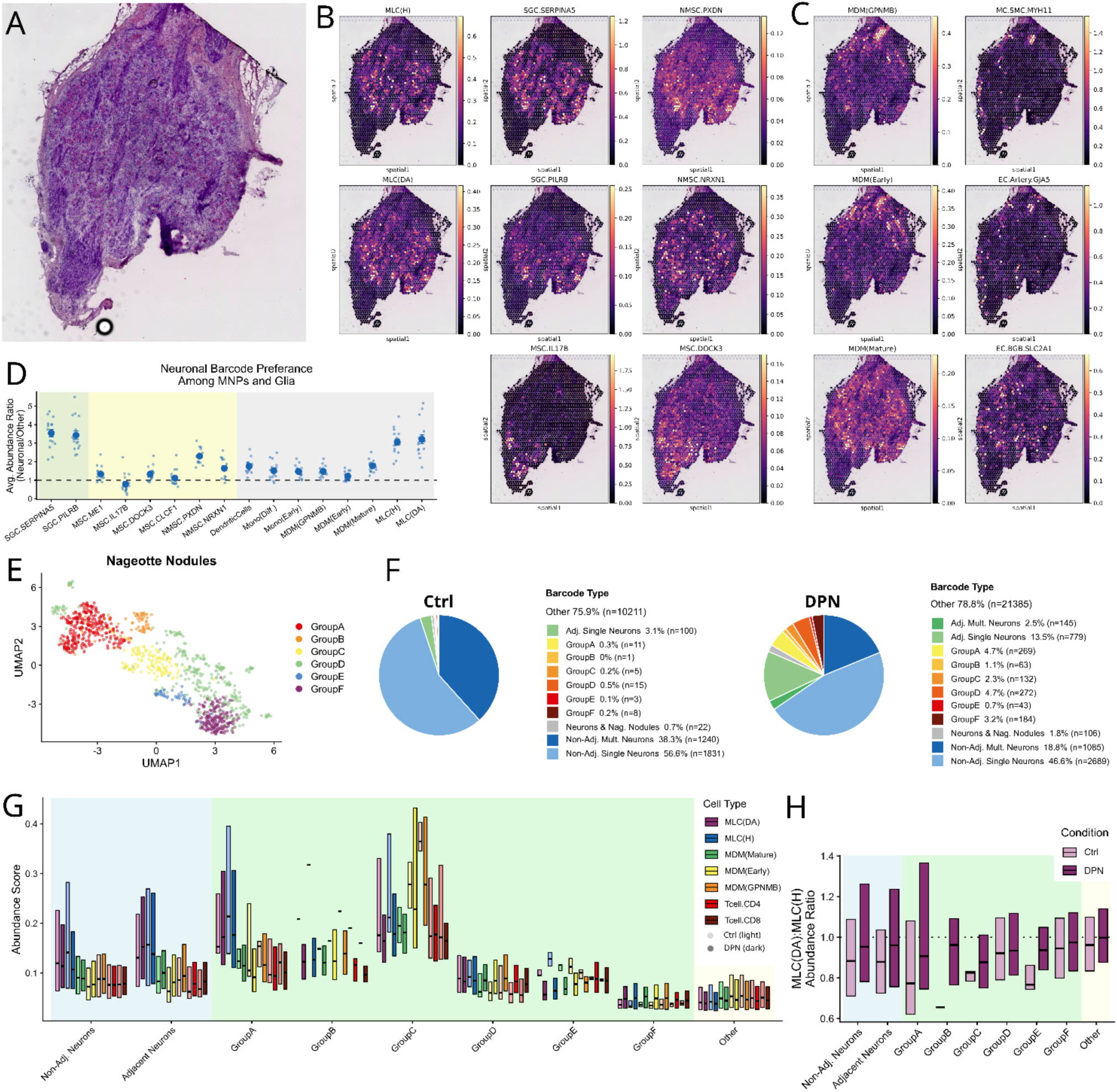
Spatial analysis revealed MLCs localize to the satellite space and exhibit changes in abundance with DPN. A) Hematoxylin and eosin stained 10x VISIUM sample from a human DRG. B) Spatial deconvolution and predicted abundance of cell types per spot revealed that MLC localization mirrored that of satellite glia populations (SGC). *PXDN*+ non-myelinating Schwann cells (NMSC) also localized to this space. Myelinating schwann cells (MSC) were enriched in regions with visible nerve tracts. C) Immature or early MDMs localized with vascular cells such as arteriolar endothelial cells and pericytes. Mature macrophages were present throughout the DRG. D) Ratio of abundance scores in manually selected neuronal barcodes to non-neuronal barcodes for satellite cell types and MNPs (data points represent average per sample; mean and SEM across samples per cell type are shown). E) Clustering of Nageotte nodule barcodes by gene expression. F) Nageotte nodules and nodule-adjacent neurons are more prevalent in human DRG samples from donors with DPN (8 samples from 3 donors) than those without diabetes (8 samples from 3 donors). Other non-neuronal barcodes are not shown but indicated in the legend. G) Nageotte nodules exhibited changes in abundance of immune cell types, with MLCs most prevalent in Group A and C nodule barcodes and MDMs most prevalent in Group C. H) Relative abundance of MLC(DA) to MLC(H) was higher in both nodule-adjacent and non-adjacent neuronal barcodes of DPN samples compared corresponding barcodes from control samples (p < 0.05).

### Glia do not express key immune cell markers in hDRG and MLCs are not glia

As MLCs have distinct gene expression patterns from MDMs and are spatially restricted to the satellite envelope^2^ unlike other macrophages, it is plausible to consider that these cells are glia. This is especially relevant given reports of satellite glial cells expressing macrophage genes. These previous studies have found that satellite glia may have antigen presentation capability ^25,62^ and express markers thought to be exclusive to immune cells such as CD45/*PTPRC*, CD14, CD68, CD83, CD11b/*ITGAM*, and Iba1/*AIF1*^20,26^. While Iba1 was shown to colocalize with cells positive for glia marker Fabp7 in rodents^20^, some of these studies did not include known markers for satellite glia and may have assumed cells in the satellite envelope were glia.

We evaluated the expression of some macrophage genes that had been observed in glia, as well as the expression of MLC markers, in all cell types of our human DRG single-nuclei dataset (Figure 2F). We found that MLC marker *WNT5A* was not expressed in glia and *ADGRG1*, while useful for differentiating MLCs from other macrophages, was highly expressed in satellite glia. The MLC markers *P2RY12*, which was utilized in previous work to validate presence of MLCs in tissue^2^, and *CD83* were also expressed by some satellite glial cells. However, many of the macrophage markers observed in satellite envelope cells, such as *PTPRC* and *ITGAM*, were not expressed by glia. Rather, they were exclusive to macrophages and other immune cells such as lymphocytes and granulocytes. Thus, we predict that *PTPRC*+/CD45+ cells in the satellite envelope are of immune and not glial lineage. While these immune cells likely include resident MLCs along with MDMs, lymphocytes, and granulocytes that enter the satellite envelope, we hypothesize that MLCs are the dominant immune cell of the satellite envelope.

To validate the immune and non-glial phenotype of MLCs, multi-parameter flow cytometry was performed on enzymatically dissociated non-neuronal cells from the lumbar level L1 DRG of a 55-year-old male organ donor. The presence of multiple expected immune and glial populations was identified, including SGCs, Schwann cells, T cells, non-MLC MNPs, and MLCs, which were defined as S100β-, FABP7-, CD45+, CD11b+, and P2Y12+ (Figure S2A). Staining for the MLC(H) marker TMEM119 showed very few positive cells in this sample; this may be attributed to the activation of these MLCs by the tissue digestion and staining procedure which is likely to result in the rapid cleaving of TMEM119^63^. A total of 50% of CD45+CD11b+ MNPs expressed the MLC marker P2Y12. In contrast, P2Y12 was expressed on only ∼15% of SGCs and negligibly on T cells and non-immune cells (Figure S2B), which is consistent with the single-nuclei analysis and confirms the relative specificity of this marker to MLCs. As a proportion of all live non-neuronal cells, MLCs constituted 5.2% of the total, which was similar in proportion to non-MLC MNPs (Figure S2C). These results confirm the presence of MLCs in the human DRG that can be distinguished from glial cells and other MNPs at the protein level.

### MLCs localize to the satellite envelope and change in abundance with diabetic painful neuropathy

Wu et al. observed that P2RY12+ cells, likely to be MLCs, reside beneath the satellite envelope^2^. However, as some satellite glia were found to express *P2RY12* in addition to MLCs (Figure 2F), we sought to assess localization of MLCs and glia whole transcriptome spatial transcriptomic data generated with the 10x Genomics Visium spatial gene expression platform. This spatial dataset consisted of 16 hDRG samples from 3 donors without diabetes and 3 donors with diabetic painful neuropathy (DPN) ^7,28^, allowing analysis of changes in MNP and glia populations with the formation of Nageotte nodules, pathological neuroma-like structures that are associated with neuronal death in hDRG^7,9^.

The hDRG single-nuclei dataset described above was used as a reference to deconvolute the spatial spots and predict localization of hDRG cell types (Figure S3 and S4) using Cell2location^64^. This spatial deconvolution tool was previously shown to perform effective differentiation of over 50 cell types, including subpopulations with subtle differences, in mouse brain^64^. Deconvolution of hDRG samples predicted localization of satellite glia to areas of the tissue slice that are visibly soma-rich and an absence of satellite glia in areas that do not have neuronal somata (Figure 3A and 3B). Similarly, MLCs are predicted to localize predominantly to soma-rich regions of the hDRG and their abundance pattern mirrors that of satellite glia (Figure 3B). MDMs are also present in soma-rich regions but, unlike MLCs, are present in other regions as well. Immature MDMs are enriched in regions with vascular cells such as arteriole endothelial cells and pericytes, while mature MDMs are spread throughout in soma-rich, axon-rich, and vascular-rich regions (Figure 3C). *NRXN1*+ non-myelinating Schwann cells seem to be distributed throughout the DRG and myelinating Schwann cells localize to axon-rich regions (Figure 3B). Interestingly, *PXDN*+ non-myelinating Schwann cells showed preference for soma-rich regions like satellite glia and MLCs (Figure 3B), suggesting that there may be non-myelinating Schwann cells resident to the satellite envelope that are different from non-myelinating Schwann cells in other regions of the hDRG. These observations were confirmed with quantification of cell abundance scores in manually annotated spatial barcodes containing somata, which showed that MLCs, satellite glia, and *PXDN*+ non-myelinating Schwann cells are enriched in neuronal barcodes compared to other barcodes (Figure 3D).

As CNS microglia respond to inflammation and can proliferate, transition, or deplete in different phases of injury resolution, we next analyzed the spatial data for changes in MLCs with DPN. Using manually annotated spatial barcodes with Nageotte nodules, largely from DPN samples, we performed unsupervised clustering by gene expression and identified 6 groups (Figure 3E and 3F). Both MLC(H) and MLC(DA) were increased in nodule-adjacent neuronal barcodes compared to non-adjacent neuronal barcodes and had the highest abundance in Group A and C nodules (Figure 3G). MLC abundance was lowest in Group F nodules along with other barcodes that did not have neuronal somata or Nageotte nodules. This pattern was observed in both non-diabetic and DPN samples and implies that MLC presence is tied to presence of the neuronal soma. Satellite glial cells and *PXDN*+ non-myelinating Schwann cells also exhibited a pattern of high abundance in Group A and C nodules and lowest abundance in Group F and other barcodes (Figure S5). As we described previously^7^, we hypothesize that the nodule clusters identified may reflect different stages of nodule development with Group A representing the earliest stage after, or perhaps during, neuronal death and Group F the latest stage. Our findings suggest that satellite resident cells proliferate around or migrate to the dying neuron and recede as formation of the Nageotte nodule progresses.

As MLCs were more abundant than MDMs among neuronal and Group A nodule barcodes (Figure 3G), they may be the first phagocytes that clear the dying neuron and may orchestrate the arrival of other immune cells. MDMs and T cells peaked in abundance in Group C nodules, and MDMs were the dominant phagocyte during this phase of nodule development. Others have also hypothesized that resident macrophages of the brain perform the first phase of phagocytosis and circulating macrophages conduct the second phase of phagocytosis after injury^9,65^. Schilling et al. even demonstrate that resident microglia contain phagocytosed neuronal debris in the first days after ischemic stroke in mice, while phagocytic monocyte-derived macrophages appeared on day 4^66^. Greenhalgh et al. have also shown that microglia are the primary cell type contacting axons and performing phagocytosis 1-3 days after spinal cord injury in mice, while monocyte-derived macrophages dominate by day 7^67^. Lastly, DPN samples showed an increase in the ratio of MLC(DA) to MLC(H) abundance in neuronal barcodes (p < 0.001 for both nodule-adjacent and non-adjacent neuronal barcodes) and Group A nodule barcodes (p = 0.05) (Figure 3H), suggesting increased MLC state transition with DPN.

### MLCs likely shift in state and intercellular signaling with diabetic neuropathy

We further evaluated changes in MLC abundance and state among sensory pathologies in the previous single-nuclei RNA-seq dataset of hDRG (Table S1). Compared to control donors without sensory pathology, MLC(H) was elevated in DRGs from donors with DPN (p = 0.01) and alcohol use disorder (p=0.02) (Figure 4A) and trending up with autoimmune disease (p=0.15). MLC(DA) was elevated in donors with DPN (p=0.002), alcohol-induced neuropathy (p=0.04), and autoimmune disease (p=0.05). Furthermore, the ratio of MLC(DA) to MLC(H) was trending up with DPN (p=0.1) and ALN (p=0.06) (Figure 4B). Increased MLCs with DPN despite neuronal loss with DPN^28^, combined with the observations that MLCs were specific to the satellite envelope of living neurons, implies that MLCs proliferate around surviving neurons or early Nageotte nodules. As seen in the spatial data (Figure 3H), these findings suggest that MLC(H) and MLC(DA) are present in hDRG in both non-diseased states and DPN, but there is a shift from MLC(H) to MLC(DA) with DPN. In human brain, Olah et al. also show that both homeostatic microglia and DAMs are present in non-diseased, neurodegenerative, and autoimmune conditions, but the relative proportions of microglial states may differ between them^68^.

**Figure 4.**
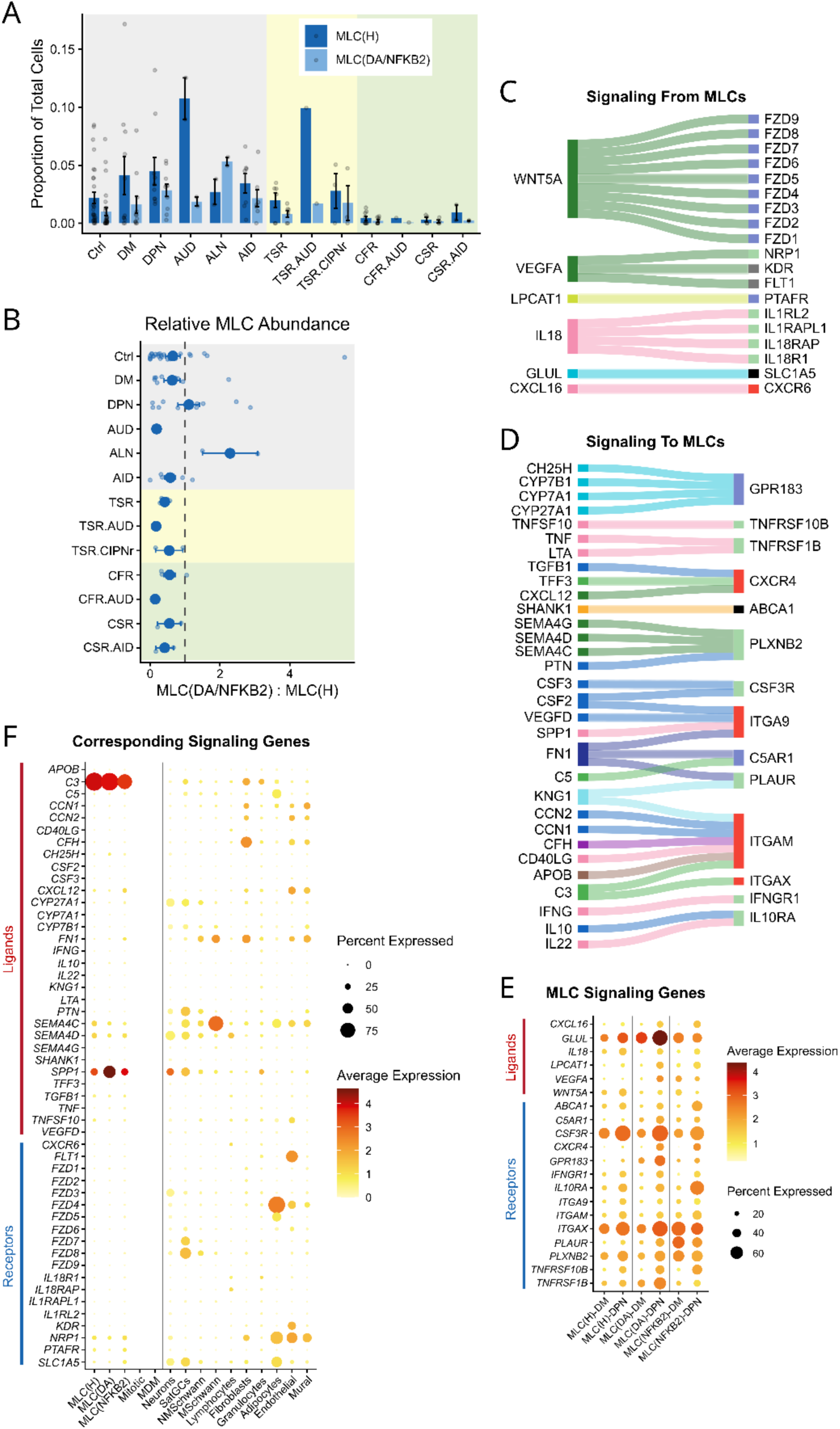
DRG pathologies change MLC abundance, activation state, and signaling mechanisms within the DRG. A) Abundance of MLCs as a percentage of all cells across pathological conditions represented in the dataset: no sensory pathology (Ctrl, n=29 donors); diabetes (DM, n=11); diabetic neuropathy (DPN, n=10); alcohol use disorder (AUD, n=2); alcohol induced neuropathy (ALN, n=2); autoimmune disease (AID, n=7); thoracic stenosis and radiculopathy (TSR, n=6); TSR with AUD (TSR.AUD, n=1); TSR with platin chemotherapy use (TSR.CIPNr, n=3); cervical fracture and radiculopathy (CFR, n=9); CFR with AUD (CFR.AUD, n=1); and cervical stenosis and radiculopathy (CSR, n=4); CSR with AID (CSR.AID, n=2); (individual data points, mean, and SEM are shown). hDRG donors included organ donors of lumbar and thoracic DRGs (gray), thoracic DRG surgical donors (yellow), and cervical DRG surgical donors (green). B) Relative abundance of MLC(DA) compared to MLC(H) was trending up with ALN and DPN; (excluding 15 donors that had no MLCs in (A)). C) Interactome of ligand genes upregulated in MLCs with DPN relative to DM (padj. < 0.05, log2FC > 1) and corresponding receptors genes expressed in the DRG. D) Receptor genes upregulated in MLCs with DPN and corresponding ligand genes expressed in the DRG. E) Expression of MLC ligand and receptor genes and F) corresponding signaling genes across DRG cell types.

To understand how MLCs may be changing functionally with DPN, we built an interactome of signaling mechanisms between MLCs and other hDRG cells that are predicted to increase with DPN (Figure 4C-F, Table S4). Using a manually curated ligand-receptor database^46^, we identified ligand and receptor genes upregulated with DPN compared to diabetes without neuropathy (DM) in any of the 3 MLC types (padj. < 0.05, log2FC > 1), along with corresponding receptor and ligand genes expressed in any of the hDRG cells. We identified 6 ligand genes upregulated with DPN in MLCs (Figure 4C). Glutamine synthase gene *GLUL* was highly expressed by all MLCs and upregulated in MLC(H) and MLC(DA) with DPN. Expression of glutamine transporter gene *SLC1A5* in neurons and satellite glia may reflect increased shuttling of glutamine from MLCs to promote glutamate production. Conversely, upregulation of *GLUL* could also reflect increased turnover of glutamate by MLCs to prevent excitotoxicity. Signaling with WNT5A, which is very specific to MLCs (Figure 2F), may be a major driver of MLC influence on other cells. Wnt5a is a neurotrophic factor for both central and peripheral nervous system neurons^69,70^, and its binding to presynaptic Fzd3 in axons promotes nerve growth^71^.

Additionally, in humans, both *WNT5A* and *FZD3* are upregulated in tension-type headache patients^72^. We observed that *FZD3* was most highly expressed in neurons and is the most highly expressed FZD in neurons (Figure 4F) and hypothesize WNT5A-FZD3 as mechanism by which MLCs could promote axonal sprouting and nociceptor sensitization in DPN. Another WNT5A receptor gene, *FZD4,* was expressed by adipocytes and vascular cells. Wnt5a and Fzd4 are downregulated in mice with increased adipogenesis^73^, suggesting WNT5A-FZD4 signaling may regulate adipocyte proliferation. Fzd4 activation in vascular cells promotes angiogenesis and impaired vascularization due to loss of Fzd4 may impair neuronal function as well^74^. Additionally, while *WNT5A* increases in MLC(H) and MLC(DA), it decreases in MLC(NFKB2) with DPN. Growth factor *VEGFA,* for which receptors were most highly expressed by endothelial and mural cells, also increases in MLC(DA) but decreases in MLC(NFKB2) with DPN. VEGFA receptor gene *NRP1* was also expressed by neurons and NRP1 is essential for VEGFA-induced nociceptor hyperactivity^75^. It may be that MLC(NFKB2), which exhibited the strongest response to inflammation with upregulation of NFKB-associated and cytokine response genes (Figure 1D), are stimulated during inflammation to inhibit axon sprouting and angiogenesis.

Additionally, we identified 14 receptor genes upregulated with DPN in MLCs (Figure 4-D). *GPR183* was upregulated in all MLCs with DPN, and is activated by oxysterols, which can be produced by CYP enzymes that were expressed by neurons and glia (Figure 4F). Oxysterol-GPR183 signaling to macrophages may serve as chemotactic cues that promote infiltration to infected tissue and stimulate production of inflammatory cytokines IL6, TNF, and IFNB^76^. GPR183 is also upregulated in spinal cord astrocytes and microglia following chronic constriction nerve injury in rats and GPR183 antagonism reverses allodynia^77^. *GPR183* was most highly expressed in MLC(DA) and could represent increased chemotaxis among MLC(DA) compared to MLC(H) and MLC(NFKB2). *IL10RA*, the primary receptor for IL10 which inhibits production of pro-inflammatory cytokines like IL6 and TNF in macrophages^78^, showed upregulation across all MLC types. This demonstrates that both pro-inflammatory and anti-inflammatory cues to MLCs may increase with DPN. However, different MLC states may differentially increase the receptors for these cues, resulting in different functional outcomes. For example, MLC(DA) expressed higher levels of *GPR183* than *IL10RA* while MLC(H) and MLC(NFKB2) expressed higher levels of *IL10RA* than *GPR183* in DPN. Additionally, *IL10RA* was most upregulated in MLC(NFKB2), further supporting the idea that this population of MLCs may play a more protective role. Furthermore, the integration of these cues could influence transition between MLC states. For example, TNF stimulates activation of canonical NFKB followed by delayed activation of non-canonical NFKB (NFKB2)^79^. Upregulation and high expression of TNF receptor gene *TNFRSF1B* (coding for TNFR2) observed in MLC(DA) may reflect induction of pro-inflammatory activity with NFKB^80^ later followed by activation of anti-inflammatory programs like IL6 inhibition with NFKB2^81^, which may also transition MLC(DA) to MLC(NFKB2). Some of the ligand genes corresponding to upregulated receptors in MLCs are also expressed by MLCs themselves, suggesting MLCs may regulate their own activity. For example, *C3* is highly expressed by MLCs and CD11 genes *ITGAM* and *ITGAX* are upregulated in MLCs with DPN. C3 fragments bind to injured cells or debris and binding to CD11b (*ITGAM*) and CD11c (*ITGAX*) promote phagocytosis. Upregulation of CD11 genes with DPN suggests increased phagocytic activity by MLCs.

### MLCs are enriched in genes involving apoptotic induction and phagocytosis

Neuronal death and removal are a hallmark of DPN pathology in hDRG^7,28^. To investigate which MNP or glia of the satellite space may be involved in promoting apoptosis and clearance of neurons or debris, we compiled 85 gene sets among Gene Ontology and Reactome terms related to these processed and grouped them into 15 modules (Figure S6). We scored the enrichment of these modules in every cell of the hDRG single-nuclei dataset and tabulated the average score per cell type for each of the modules (Figure 5A). Most phagocytosis processes were enriched in MNP cell types while some processes such as lipid and amyloid clearance, were enriched in satellite glial cells. MLCs were also among cell types implicated in cytotoxicity, apoptotic induction, and phagocytosis of apoptotic cells. Interestingly, MLC(NFKB2) were also enriched in processes that may inhibit apoptosis. As most phagocytosis processes were enriched in MNPs overall, we evaluated which genes from these modules were enriched in MLCs compared to other MNPs (padj. < 0.05 and log2FC >1) (Figure 5B). We observed that *C3* and *ITGAX* are more highly expressed in MLCs compared to MDMs. However, most of the genes enriched in MLCs were expressed by other MNPs as well. This suggests that both MNPs and other macrophages may perform phagocytosis. As MLCs were found to be the major MNP of the peri-neuronal space and MDMs were found to peak later during the process of Nageotte nodule development (Figure 3H), we hypothesize that MLCs play an important role in phagocytosis immediately after irreversible neuronal damage and that other macrophages may be the dominant phagocytes in later stages of debris clearing.

**Figure 5.**
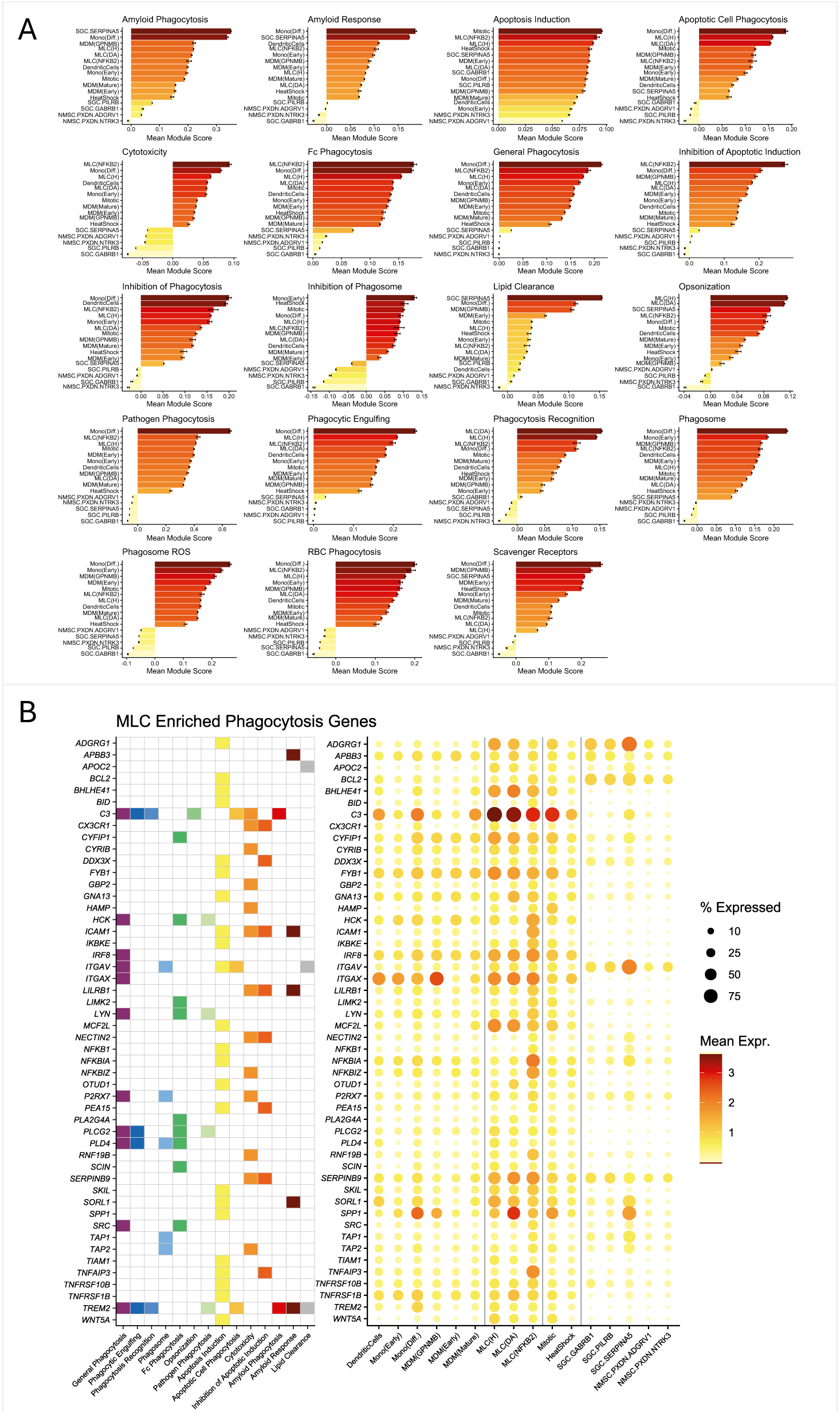
MLCs are enriched in genes associated with phagocytosis mechanisms. A) MNPs and glial cells of the satellite space were scored by expression of genes in each of 15 modules covering different phagocytosis or cell death mechanisms from 85 Gene Ontology and Reactome terms (mean and SEM shown per cell type). Most mechanisms were enriched in MNP cell types with some mechanisms such as clearance of lipids and amyloid enriched in satellite glia. B) Several genes across the modules showed enrichment in MLCs relative to MDMs (padj. < 0.05, log2FC > 1).

### MLC markers are detected in other PNS structures

We examined MNPs in published single-nuclei datasets from other peripheral ganglia and found clusters of cells expressing microglia and MLC markers in each case. In human sympathetic chain ganglia^36^, we identified a group of MLCs enriched in *ADGRG1* and *WNT5A,* with subpopulations enriched in MLC(H) marker *TMEM119*, MLC(DA) marker *CD83*, and mitotic marker *MKI67* (Figure 6A). We also identified MDMs enriched in *MRC1*, monocytes enriched in *CD300E*, and dendritic cells enriched in *CLEC10A*. Similarly, in human trigeminal ganglia^35^, we identified a *ADGRG1+WNT5A+* MLCs with *TMEM119+* and *CD83+* subpopulations, along with *MRC1+* MDMs and a *ADGRG1+WNT5A+MRC1-*mitotic cluster (Figure 6B). As there are no published datasets of human nodose ganglia, we examined a barcoded spatial dataset of human nodose^34^ for MLC markers. MLC markers were detected in manually annotated neuronal barcodes, with *CX3CR1* the most sensitive marker for MLC(H) and *CCL3* the most sensitive marker for MLC(DA) (Figure 6C). MLC markers were enriched in neuronal barcodes relative to other barcodes (Figure 6D). Moreover, the ratio of *CCL3+* neuronal barcodes to *CX3CR1+* neuronal barcodes was increased in donors 38 and 26, who had a history of diabetes. Nodose tissue from donor 26, who had the highest ratio of *CCL3+* to *CX3XR1+* neuronal barcodes, also exhibited abundant Nageotte nodules (Figure 6E). These findings suggest that MLCs are present throughout PNS ganglia. It may be that the MLCs of other ganglia are different from MLCs of the DRG and their similarity is just tied to expression of microglia markers by all resident macrophages of the nervous system. However, enrichment of MLC markers in neuronal barcodes in the nodose ganglia suggests that, like hDRG MLCs, MLCs of other ganglia reside in the satellite envelope. Additionally, increase in MLC(DA) marker genes relative to MLC(H) marker genes in neuronal barcodes of likely diseased Nodose parallels MLC dynamics seen in hDRG (Figure 3I).

**Figure 6.**
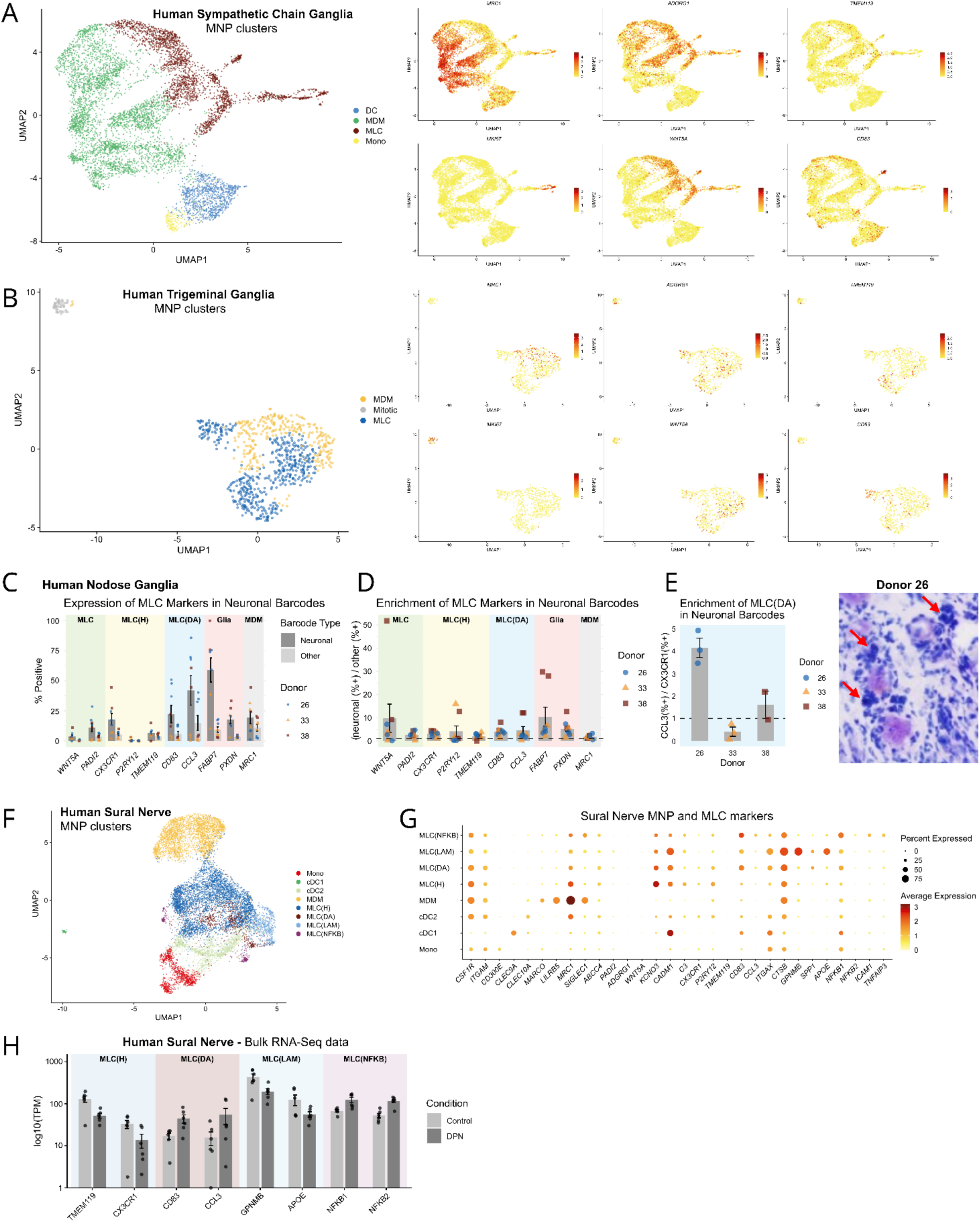
MLCs are present in other peripheral ganglia and exhibit changes with DPN in sural nerves. Peripheral nervous system MNPs likely consist of MDMs and MLCs as observed in single-cell datasets of A) human trigeminal ganglia, B) human sympathetic chain ganglia, and F) human sural nerve. MDM cells were enriched in *MRC1* across these datasets, while *WNT5A* and *ADGRG1* were markers for MLCs in ganglia but not sural nerve, where *KCNQ3* was a more sensitive marker. Subpopulations enriched in MLC(H) markers *TMEM119* or *P2RY12,* and MLC(DA) marker *CD83* were also observed. H) A bulk RNA-seq dataset of human sural nerve showed decrease of MLC(H) markers and increase of MLC(DA) markers with DPN. C) A 10x Visium dataset of human nodose revealed expression of MLC markers in neuronal barcodes, with *CX3CR1* and *CD83* as the most highly expressed markers for MLC(H) and MLC(DA), respectively. D) Some MLC markers were enriched in neuronal barcodes compared to other barcodes, as were the SGC marker *FABP7* and satellite Schwann cell marker *PXDN*. E) Comparison of the proportion of neuronal barcodes positive for *CCL3* and *CX3CR1* suggested increased abundance of MLC(DA) relative to MLC(H) in donors 26 and 38, who had a history of diabetes (left). Nodose samples from donor 26 exhibited Nageotte nodules throughout the ganglion (right).

There are also reports of endoneurial macrophages that express microglia genes^82^ and CX3CR1+ macrophages that respond to injury in peripheral nerves^33^. We examined all MNPs from a single-nuclei dataset of human sural nerve by Heming et al.^33^ and identified clusters of dendritic cells, monocytes, MDMs, and MLCs (Figure 6F). While MLCs did not robustly express many of the MLC markers from peripheral ganglia, such as *WNT5A*, *ADGRG1, PADI2,* they were enriched in *C3, CADM1,* and *KCNQ3* (Figure 6G). Additionally, we found MLC subpopulations enriched in MLC(H) marker *P2RY12*, MLC(DA) marker *CD83,* and NFKB genes. We also identified a large cluster of MNPs enriched in lipid processing genes *GPNMB*, *SPP1*, and *APOE* which were termed lipid-associated macrophages (LAM) by Heming et al. We predict that these cells are also MLCs because they express *C3, CADM1,* and *KCNQ3,* and express *MRC1* at low levels like the other MLC groups in the sural nerve. Analysis of bulk transcriptomic data from non-diseased and DPN human sural nerves^32^ showed that expression of MLC markers changes with DPN (Figure 6H). Markers for MLC(H) *TMEM119* and *CX3CR1* decrease with DPN, as do markers for MLC(LAM) *GPNMB* and *APOE*. MLC(DA) markers *CD83* and *CCL3* increase with DPN. MLC(NFKB) markers *NFKB1* and *NFKB2* also increase with DPN, but these genes may be expressed by other cell types as well. Alternate MLC marker expression and a unique MLC subpopulation in sural nerve suggests that nerve MLCs may be different from MLCs of peripheral ganglia. However, MLCs of the sural nerve and the ganglia studied here also shared microglia markers, which may be common for all resident macrophages of the nervous system. All PNS datasets also exhibited homeostatic and disease-associated MLC, and there were similar changes with DPN in human DRG, nodose, and sural nerve.

## Discussion

Microglia-like cells are likely the resident tissue macrophages of the peripheral nervous system. We used single-nuclei and spatial datasets to confirm the presence of MLCs in hDRG and have substantially expanded on their transcriptomic profile, including characteristics that distinguish them from other MNPs and glia. We also predicted MLC functional traits and how they signal with neurons. We identified homeostatic and disease-associated MLC subpopulations that may reflect cell state transitions, and we explored how cellular activities may change across these states. We provided further evidence that MLCs reside in the satellite envelope as described by Wu et al. ^2^, but include a whole transcriptome approach that evaluates localization of all hDRG cell types simultaneously using several markers for each cell type. Additionally, we show that localization of MLCs and other MNPs to the peri-neuronal space changes with DPN, in a manner that suggests MLCs drive phagocytosis in the earliest stages of Nageotte nodule formation and that MDMs are the dominant phagocytes in later stages. We hypothesized the specific signaling mechanisms by which MLCs may drive and respond to the activities of other hDRG cells in DPN. Lastly, we identified likely MLCs in published datasets of other peripheral ganglia and sural nerve. We conclude that MLCs are likely present throughout the peripheral nervous system and changes seen with peripheral neuropathy may reflect their complex role in both promoting and protecting against inflammation.

### MLCs: a new focus on a commonly observed but mysterious cell type

While the concept of MLCs in the DRG may seem new, it is very likely that others have observed them in sensory ganglia in studies going back decades^2–24^ (Table 1). It has been known since at least the early 1990s that there are resident phagocytic cells around neuronal somata in sensory ganglia^83,84^ but there have been different interpretations of whether they are marrow-derived macrophages or specialized macrophages resident to the peripheral ganglia. Smith et al. showed that the number of satellite cells in mouse DRG increases after sciatic nerve injury and that some of them were positive for mononuclear phagocyte markers indicating macrophage lineage^85^. Esiri et al. studied human sensory ganglia (TG and DRG) and sympathetic ganglia (superior cervical, thoracic, and celiac) and described satellite macrophages that are present even in the absence of disease and have staining characteristics different from interstitial macrophages of the ganglia^86^. Fiori et al. observed satellite macrophages in chick ciliary ganglia that resemble CNS microglia, and were the first to term them ‘microglia-like cells’. They also describe ultrastructural characteristics of these cells as having elongated, electron-dense cell bodies and lack of basal lamina. Scaravilli et al. described a population of satellite cells in human DRGs in the same manner and observed that they are not present in Nageotte nodules or in rats. They further described them as having ‘cigar-shaped’ nuclei, embedded within the processes of other satellite cells, and sometimes located directly on the neuronal somata.

### Differentiating MLCs from other cell types

Given their localization to the same space in the satellite envelope, it could seem that MLCs are satellite glia. With the perceived absence of microglia or other resident immune cells specific to peripheral ganglia, it has been thought that satellite glia may help fill the role that microglia play in the central nervous system^26^ and some findings report the expression of immune cell markers in satellite glia, such as CD45/*PTPRC*, CD11b/*ITGAM*, and CD11c/*ITGAX*^26,27^. However, these cells may have been assumed to be SGCs due to their morphology and peri-soma localization. We found that SGCs do not express many key immune markers like *PTPRC, ITGAM,* and *ITGAX*, and most hDRG satellite cells enriched in these genes are likely MLCs. As the MLCs are negative for glia markers, share transcriptomic markers with MDMs, and cluster with MDMs in our single-nuclei data as well as previous studies, they are usually labeled as MDMs in single-cell datasets of sensory ganglia^30^. We provided markers, such as *WNT5A*, that can differentiate MLCs from MNPs and all other hDRG cell types. We predict that WNT5A expression by MLCs can influence activity in several hDRG cell types by signaling to FZD receptors and that it may allow MLCs to promote axon sprouting and angiogenesis in DPN. We also predicted other functional differences between MLCs and MDMs, including increased cell adhesion with neurons, signaling with axonal growth cues, and regulation of neurotransmitter dynamics. We provided further support for MLCs being resident tissue macrophages in hDRG with trajectory analysis in post-natal DRGs that showed MLCs likely do not arise from monocytes and with analysis of embryonic human DRGs that showed MLC precursors are found only during development, as is seen with resident tissue macrophages of other tissues. Lastly, we showed that MLCs localize to the peri-neuronal space while immature MDMs localize to vascular-rich regions and mature MDMs are spread throughout the hDRG.

### MLCs in DPN pathogenesis

We found that MLCs exhibit changes in abundance, localization, cell state, and signaling mechanisms with DPN. We show that in both hDRG and human sural nerve, the relative abundance of MLC(DA) to MLC(H) increased with DPN. This may represent a shift in cell state to MLC(DA) or increased proliferation of these cells, which upregulated *GPR183* in DPN, and may promote inflammatory cytokines like IL6 and nociceptor sensitization. Additionally, we postulate that MLC(NFKB2), while exhibiting a transcriptomic profile of response to inflammation, may actually play a protective role. This MLC group showed significant upregulation of *IL10RA*, which can inhibit production of inflammatory cytokines including IL6^78^. Additionally, TNF stimulation of MLC(DA) may induce NFKB2 promoting transition to the MLC(NFKB2) state.

We also observed that MLCs increased in neuronal spots that were adjacent to Nageotte nodules compared to non-adjacent neuronal barcodes and increased further in what may be the earliest stage of Nageotte nodule formation. Up to that point, MLCs were the dominant MNP of the peri-neuronal space, and then exhibited significant decline in other nodule groups, suggesting that neuronal presence is necessary for MLCs. Other MNPs increased in what may be later stages of nodule development, suggesting that MLCs are more abundant in the earliest stages of neuronal death while other phagocytes infiltrate in a later stage. While other satellite cells, including satellite glia and *PXDN*+ non-myelinating Schwann cells, displayed the same abundance patterns as MLCs with increase in nodule-adjacent neurons and further increase in a group of nodules that may be in the earliest stage, we predict that MLCs likely play a larger role in phagocytosis of dying neurons because phagocytosis programs are highly enriched in MNPs compared to glia. We also observed that MLCs express genes implicated in cytotoxicity and apoptotic induction, and may play role in promoting neuronal death in DPN. MLC(NFKB2) was also enriched in genes that may inhibit apoptosis, further indicating that MLCs could play a protective role in DPN.

### Limitations and future directions

The computational analysis of MLCs presented here provides novel insight into MLCs of the hDRG and their role in inflammation, nociceptor sensitization, axonal sprouting, and neuronal death. It also provides evidence for MLCs as the resident tissue macrophage of the hDRG and their presence throughout the PNS. However, future studies that examine MLCs in tissue using specific markers would help validate these findings as sequencing analysis can have limitations, although our findings are based on multiple, independent methods. An example of such a limitation is that MLC(H) increased in number with DPN in the hDRG single-nuclei dataset, but predicted abundance of MLC(H) in neuronal barcodes was not increased with DPN in the spatial hDRG data, and MLC(H) markers decreased with DPN in analysis of bulk RNA-seq data from human sural nerves. This suggests that sequencing techniques may bias detection of MLCs and in-situ analysis of MLC subpopulations may help validate their abundance changes with peripheral neuropathy. Additionally, functional experiments of neuronal response to MLC signaling molecules like *WNT5A* or *VEGFA* would help validate their predicted role in neuronal sprouting and sensitization. With this work, we aim to bring attention to MLCs as a key player in regulating neuronal function and peripheral neuropathy. We hope that it will open a new chapter in investigating and targeting MLC functions for pathologies across the PNS.

## Methods

### Ethics approval for human DRG procurement

Human tissue procurement procedures were approved by the Institutional Review Board at the University of Texas at Dallas (MR-15-237). The Southwest Transplant Alliance (STA) obtained informed consent for research tissue donation from first-person consent or from the donor’s legal next of kin. Policies for donor screening and consent are those established by the United Network for Organ Sharing (UNOS). STA follows the standards and procedures established by the US Centers for Disease Control (CDC) and are inspected biannually by the Department of Health and Human Services (DHHS). The distribution of donor medical information is in compliance with HIPAA regulations to protect donor privacy.

### Analysis of human DRG single-nuclei RNA-sequencing data

The human DRG single-nuclei dataset consisted of 92 samples prepared with the 10x Genomics Chromium Single Cell Gene Expression Flex assay^28^, of which 80 were derived from published datasets^7,28–31^ and 12 had not been previously analyzed (Table S1). Raw sequencing files were aligned (using reference genome GRCh38) with 10x CellRanger to generate count matrices per sample, which were then processed with ambient RNA correction tool SoupX^87^ and doublet correction tool scDblFinder^88^. The hDRG cells were clustered with the Seurat v5 workflow^89^ using default parameters and harmony integration^90^. Cell types were identified by evaluating expression of known marker genes^30^ as well as top enriched genes per cluster. High-level cell types were further subclustered in a similar manner, and subclusters expressing markers exclusive to other cell types were considered to be contaminated and removed. These methods, along with specific analyses described below, were completed with R version 4.3.3.

1. <u>MLC Subtype Analysis.</u> Differentially expressed genes (DEGs) for each MLC subtype compared to other MLCs were identified using Seurat FindMarkers() with Wilcoxon rank-sum test followed with and hits (padj. < 0.1, expression in greater than 30% of the given MLC type, and expression in less than 50% of other MLCs) were assessed again with MAST using sample origin as a latent variable to control for batch effect and inter-individual variability. Enriched genes per MLC type (padj. < 0.05 after MAST) were compared with Gene Ontology Biological Process^39^ terms with the enrichR^40^ package and the top 25 enriched terms (padj. < 0.05) were selected. These terms were grouped into high level categories and expression of constituent genes was visualized using Seurat DotPlot() and gene annotations with R package ggplot2.
2. <u>Trajectory Analysis of MNPs</u>. The Slingshot algorithm^44^ was applied to the UMAP of hDRG MNPs, after exclusion of the Mitotic and Heat Shock groups that were likely heterogeneous and expressed markers from multiple MNP types. Slingshot was run with omega = TRUE. dist.method = “simple”, and default settings for other parameters.
3. <u>Interactome analysis of MLC and MDM signaling to neurons</u>. DEGs between MLCs and MDMs MDM(Early), MDM(Mature), MDM(GPNMB)) were identified using Seurat FindMarkers() with the Wilcoxon and MAST tests as was done with MLC subtype analysis. Pseudbobulked neuronal gene expression transformed with Seurat NormalizeData() was used to identify genes expressed in neurons, along with highly expressed neuronal genes (normalized expression > 0.1). MLC DEGs and neuronal genes were intersected with a curated database^46^ of ligand-receptor pairs to generate an interactome of potential signaling with neurons that is enriched in MLCs compared to MDMs. Similarly, an interactome was generated for MDM-neuronal signaling, and top interactions (with neuronal gene expression > 0.1) were visualized with SankeyMATIC (sankeymatic.com). WikiPathways^61^ terms enriched among MLC signaling genes were identified with Enrichr, along with terms enriched among corresponding neuronal genes. The padj. values and the genes associated with the pathway terms were visualized with ggplot2.
4. <u>MLC abundance across conditions.</u> Cell counts were tabulated per sample and per MLC subtype within each sample. Samples from the same donor were combined, and counts for MLC(DA) and MLC(NFKB2) were also combined. MLC(H) and MLC(DA/NFKB2) proportions as a fraction of total cells and the MLC(DA/NFKB2):MLC(H) ratio per donor were visualized with ggplot2, along with mean and SEM. Pairwise Wilcoxon rank-sum tests using the rstatix package were performed to assess differences in MLC abundance and the MLC(DA/NFKB2):MLC(H) ratio.
5. <u>Interactome analysis of MLC signaling changes with DPN.</u> Genes upregulated for each MLC type in DPN samples compared to non-neuropathic diabetes samples were identified using Seurat FindMarkers() with the Wilcoxon and MAST tests as described above. Ligand genes among the DEGs were identified using the same curated interaction database^46^, along with corresponding receptors, generating interactome of signaling from MLCs that may be upregulated with DPN. Similarly, signaling mechanisms to MLCs in DPN were identified from MLC DEGs that correspond to receptors. Interactomes were visualized with SankeyMATIC and expression of ligand and receptors genes were visualized with Seurat DotPlot().
6. <u>Analysis of phagocytosis gene sets</u>. The MSigDB gene set database^91^ was accessed with the msigdbr R package to identify 85 gene sets implicated in phagocytosis and apoptosis processes. These gene sets were manually reviewed and combined into 15 groups, or modules. All hDRG cells were scored for each module using Seurat AddModuleScore(). Module scores were visualized as bar plots (mean ± SEM) and as z-scored dot plots for all MNP and satellite glia cell types. Differential expression of phagocytosis genes between each MLC subtype and pooled non-MLC MNPs was tested using Seurat FindAllMarkers() with Wilcoxon rank-sum test. Genes enriched in any MLC type (padj. < 0.001, log2 fold-change > 1, expression in at least 25% of cells, and expression in MDMs less than 50%), and were visualized Seurat DotPlot() and categories were annotated with gglpot2.

### Analysis of human DRG spatial RNA-sequencing Data

We utilized a published 10x Visium hDRG dataset^7,28^ consisting of 16 samples from 3 donors with DPN and 3 donors with no history of diabetes. This dataset included annotations of barcodes containing Nageotte nodules, neurons adjacent to nodules, neurons not adjacent to nodules, and other barcodes. The hDRG single-nuc dataset’s count matrix and cell type labels were used the create a signature matrix and deconvolute the spatial data using Cell2location^64^. The q05 posterior distribution estimates were used as the cell type abundance scores per spatial spot and visualized as a representative slide as instructed in the Cell2location tutorial (cell2location.readthedocs.io). Observations on cell type abundance patterns across hDRG samples were made, and each cell type’s enrichment to neurons or the satellite envelope was quantified by dividing average abundance score in neuronal barcodes by average abundance score in other barcodes per sample.

Spatial barcodes with Nageotte nodules were clustered based on gene expression using the Seurat workflow with default parameters and 6 groups were identified. The median and IQR abundance scores of MLCs, MDMs, and T cells were visualized for each barcode type using ggplot2. The ratio of MLC(DA) abundance score to MLC(H) abundance score was also evaluated per barcode type, and changes in the ratio with DPN for each barcode type were determined with Wilcoxon rank-sum tests using the rstatix package. For each cell type, enrichment across barcode types was evaluated by scaling the average abundance score per barcode type with z-score. For each cell type, the scaled abundance scores and percent of barcodes in each barcode type that had abundance score greater than z=1 were visualized on a dot plot with ggplot2.

### Analysis of other published CNS and PNS RNA-sequencing datasets

Other published single cell, spatial, and bulk RNA-sequencing datasets were analyzed to compare hDRG MLCs with microglia, investigate MLC precursors, and determine whether MLCs are present in other PNS structures.

1. <u>Comparison with human brain microglia.</u> MNPs from hDRG were mapped onto a single cell dataset of human brain MNPs (including 12 microglia clusters and 1 MDM cluster) by Sun et al.^41^ using Seurat function MapQuery(). This process simultaneously determined which brain MNP type each hDRG MNP cell was most similar to and plotted the hDRG MNPs on the brain MNP UMAP. hDRG cells with a prediction score of <0.5 were considered unassigned. Differential expression of microglial genes known to be enriched in homeostatic or disease-associated microglia from rodent studies^42^ was evaluated with Seurat FindMarkers() using the Wilcoxon test. These genes were evaluated for enrichment in two homeostatic microglia compared to other microglia clusters, and between the MLCs clusters relative to each other. Log10 fold-changes and padj. values were visualized with ggplot2.
2. <u>Analysis of embryonic hDRG.</u> A single nuclei dataset of hDRG cells during embryonic development (gestational week 7-21) by Lu et al.^45^ contained a cluster of macrophages, which were subclustered using the Seurat workflow with default parameters. Expression of MLC and MDM markers across macrophage clusters was visualized with Seurat DotPlot(). Of the 2 clusters that expressed MLC markers, one also expressed MDM markers as is predicted in MLC precursors. Cell counts for each MNP type per gestational week were tabulated and visualized as a stacked bar plot with ggplot2.
3. <u>Human trigeminal and sympathetic ganglia.</u> MNPs from a single nuclei dataset of human trigeminal ganglia by Yang et al.^35^ and a human sympathetic chain ganglia dataset by Yang et al.^36^ were clustered using the Seurat workflow with default parameters. Seurat FeaturePlot() was used to visualize expression of general MLC markers *ADGRG1* and *WNT5A*, MLC(H) marker *TMEM119*, MLC(DA) marker *CD83*, MDM marker *MDM1*, and mitotic marker *MKI67* among MNP clusters.
4. <u>Human sural nerve.</u> MNPs from a single nuclei dataset by Heming et al.^33^ of sural nerve biopsies, including samples from donors with peripheral neuropathy were clustered using the Seurat workflow with default parameters. Seurat DotPlot() was used to visualize expression of MLC and MDM markers, as well as markers for lipid-associated macrophages described by Heming et al. A bulk dataset by Tavares-Ferreira et al.^32^ of sural nerve biopsies from donors with DPN and no nerve disease was also analyzed for expression of MLC markers. Bar plots showing individual samples, mean, and SEM were generated with ggplot2.
5. <u>Human nodose ganglia.</u> The only published dataset of human nodose is a 10x Visium dataset with 8 samples, including 6 samples from donors with diabetes, and manual annotation of barcodes containing neurons^34^. Expression of MLC, MDM, and glial markers in neuronal and other barcodes was visualized with data points reflecting percentage of barcodes in a sample expressing the marker and bars reflecting mean ± SEM across samples. Enrichment of markers in neurons was also evaluated by dividing the percentage of neuronal barcodes expressing the marker by the percentage of other barcodes expressing the marker per sample. We observed that all 3 samples from one of the donors with diabetes exhibited abundant Nageotte nodules and evaluated for donor differences in the ratio of neuronal barcodes expressing MLC(DA) marker *CCL3* to neuronal barcodes expressing MLC(H) marker *CX3CR1.* Sample values, mean, and SEM were visualized in a bar plot with ggplot2.

### Flow cytometry analysis of hDRG cells

1. <u>Human dorsal root ganglion tissue digestion.</u> A lumbar (L1) level DRG from a 55-year-old male donor was recovered post-mortem as previously described^92^ and stored in artificial cerebrospinal fluid on ice for immediate transport from the operating room to the laboratory. Upon arrival at the laboratory, the DRG was trimmed of extraneous nerve and adipose tissue, cut into 1 mm pieces, and placed in 5 mL of 2 mg/mL collagenase type 4 (Worthington LS004188) with 4 mg/mL DNase I (Worthington LS002139) in Hanks’ balanced salt solution with calcium and magnesium (Gibco 24020-117) in a shaking water bath at 37°C. After 3 hours, mechanical triturations with a fire-polished glass pipette were performed every 30 min. After 5 total hours of digestion, cells were placed through a 70 µm nylon mesh strainer, centrifuged at 300 *g* for 5 min at 4°C, and resuspended in 900 µL of 0.5% bovine serum albumin (BSA) in phosphate-buffered saline (PBS, pH 7.4). Cell counts were estimated using a Neubauer hemocytometer. Myelin depletion was performed on the resulting cell suspension. Myelin removal beads (Miltenyi 130-096-733, 100 µL) were added to the cell suspension and incubated for 15 min on ice. The reaction was stopped by adding 2 mL of PBS, centrifugation at 300 *g* for 5 min at RT, and aspiration of the supernatant. The pellet was resuspended in 1 mL of 0.5% BSA in PBS and passed through a pre-washed LS column (Miltenyi 130-042-401) mounted in a MidiMACS separator (Miltentyi 130-042-302). The column was washed with a further 2 mL of 0.5% BSA in PBS and the elute was centrifuged at 300 *g* for 5 min at RT.
2. <u>Flow cytometry staining.</u> All centrifugation/wash steps were performed at 300 *g* for 5 min at 4°C prior to fixation and at 800 g for 5 min at 4°C after fixation. Digested cells were resuspended in 200 µL PBS. Viability staining was performed by adding 2 µL of Zombie-UV (Biolegend 423107) and incubating for 15 min at RT. Cells were washed with flow cytometry staining buffer (eBioscience 00-4222026) and resuspended in 200 µL of 1:40 Fc block (Biolegend 422302) in staining buffer and incubated for 10 min on ice. The primary antibody cocktail for surface markers was added without washing and incubated for 20 min on ice. Cells were washed with staining buffer and then fixed in Fix/permeabilization buffer (Biolegend 426803) for 20 min at RT. Cells were washed in staining buffer containing 1:1000 tandem stabilizer (Biolegend 421802) and left as a pellet at 4°C overnight. The next day, cells were washed in permeabilization buffer and resuspended in 100 µL of intracellular primary antibody mix and incubated for 20 min on ice, followed by a permeabilization buffer wash and resupension in intracellular secondary reagent mix for 20 min on ice. After one wash each in permeabilization buffer and staining buffer, cells were resuspended in 500 µL of staining buffer and kept on ice. Single-stained compensation controls for each antibody were performed using UltraComp eBeads (Invitrogen 01-3333-42). The live-dead compensation control used ArC amine-reactive beads (Invitrogen A10346). Data was acquired on a five-laser BD Fortessa and the compensated data was gated in FlowJo v10.10. The resulting data analysis and visualization was performed in *R* v4.6.0 using RStudio.

## Resource Availability

All data and code used in this manuscript will be made publicly available on SPARC.science at DOI: (10.26275/f8wk-3m8t), including input files and code used to analyze previously published datasets from 1) postnatal human DRGs^7,28–31^, 2) human brain MNPs^41^, 3) human embryonic DRGs^45^, 4) human trigeminal ganglia^35^, 5) human sympathetic ganglia^36^, 6) human nodose ganglia^34^, and 7) human sural nerve^32,33^.

## Acknowledgements

The authors thank the organ donors and their families for their enduring gift, as well as our partners at Southwest Transplant Alliance without whom this research would not be possible. The authors also thank the Genome Center at The University of Texas at Dallas for the services to support our research. We gratefully acknowledge the members of the UTD recovery team: Stephanie Shiers, Katherin Gabirel, Keerthana Natarajan, Sera Nakisli, Joseph Lesnak, Moeno Kume, Lucy He, Marisol Mancilla Moreno, Urzula Enzastiga, Asta Arendt-Tranholm, Jayden O’Brien, Khadijah Mazhar, Kathleen Domalogdog, Ryan McKee, Nishka Kuttanna, Zulmary Manjarres Farias, Alejandro Otero Pedraza, Pegah Haghighi, Andi Wangzhou, Seph Palomino, Dhananjay Naik, Salma Ashshareef, and Ifunanya Okolie. We acknowledge the equipment, resources, and technical expertise of Jacob Henderson and the University of Texas at Dallas Flow Cytometry Core Research Facility.

## Author contributions

K.M.: Conceptualization, Data curation, Formal analysis, Investigation, Methodology, Project administration, Software, Supervision, Validation, Visualization, Writing – original draft, Writing – review & editing. J.O.: Formal analysis, Investigation, Methodology, Validation, Visualization, Resources, Writing – original draft, Writing – review & editing. M.W.: Formal analysis, Visualization, Writing – review & editing. H.S.: Formal analysis, Visualization, Writing – review & editing. A.W.: Formal analysis, Methodology, Writing – review & editing. V.P.: Formal analysis, Investigation, Writing – review & editing. C.M.: Formal analysis, Investigation, Writing – review & editing. N.A.: Writing – original draft, Writing – review & editing. M.M.M.: Formal analysis, Investigation, Writing – review & editing. I.S.: Data curation, Investigation, Resources, Writing – review & editing. D.T.: Data curation, Investigation, Resources, Writing – review & editing. T.P.: Conceptualization, Funding acquisition, Methodology, Project administration, Resources, Supervision, Writing – original draft, Writing – review & editing.

