## Supplementary Figures for "Satellite microglia-like cells in human dorsal root ganglia and changes with diabetic neuropathy"

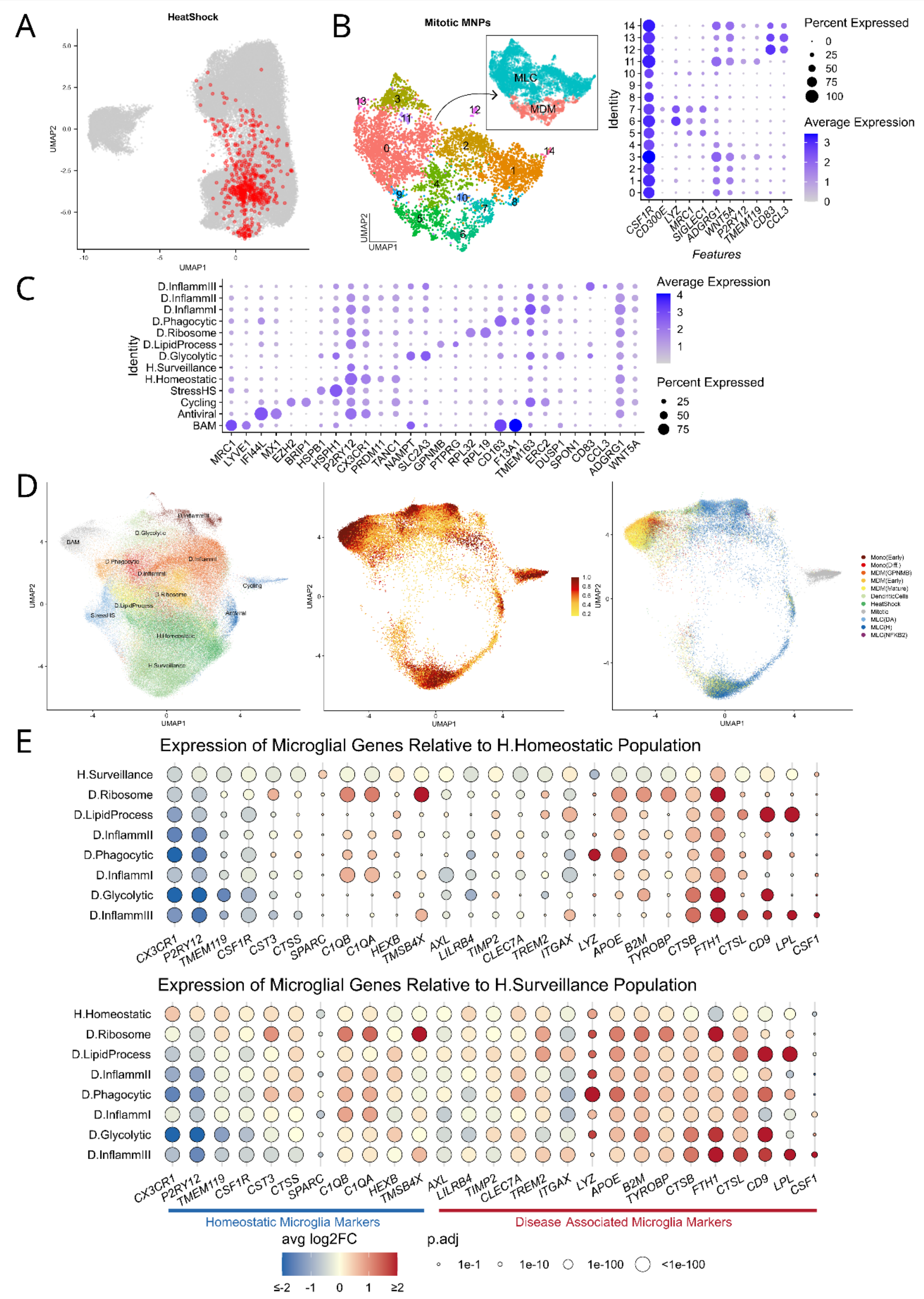

**Figure S1. Additional analysis of MNPs from hDRG and human brain**
A) Heat shock cells are distributed throughout the UMAP of hDRG MNPs and likely represent multiple MNP types in this state. B) Subclusters of mitotic MNPs were largely *ADGRG1+WNT5A+* MLCs - including *CD83+* and *TMEM119+* clusters - and *MRC1+SIGLEC1+* MDMs. C) Human brain MNPs and their markers as described by Sun et al., along with expression of MLC markers *ADGRG1* and *WNT5A*. D) UMAP of brain MNPs (left), hDRG MNPs mapped onto the UMAP (right) along with prediction scores (middle). E) Enrichment of HM and DAM markers known from rodent studies in Homeostatic microglia (top) and Surveillance microglia (bottom) compared to all other microglia clusters (excluding altered states such as stressed, antiviral, and cycling).

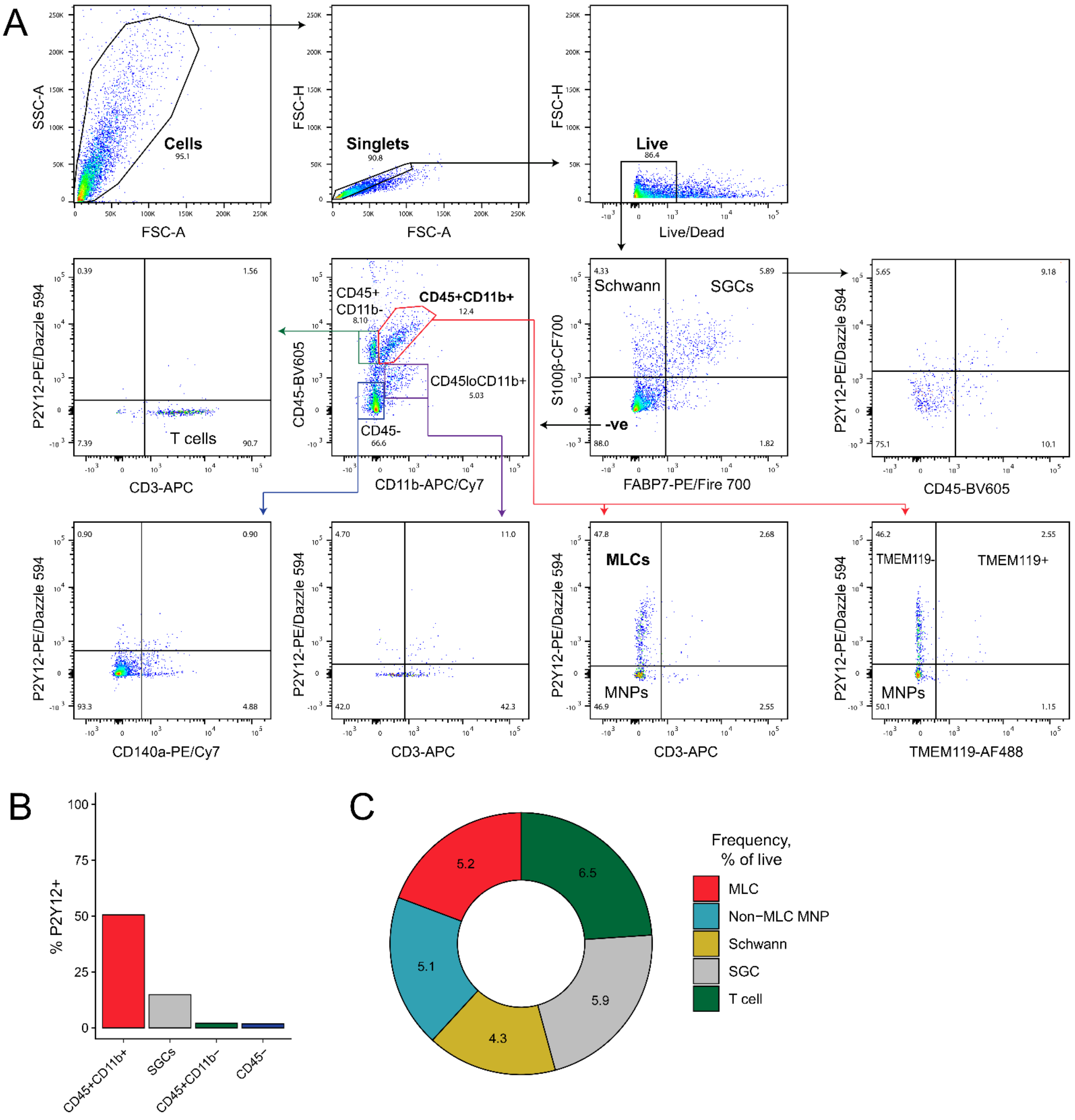


**Figure S2. Isolating MLCs in hDRG with flow cytometry**
Flow cytometry on non-neuronal cells isolated from a human dorsal root ganglion confirms the presence of a P2Y12+ cell population of MLCs that is distinct from other mononuclear phagocytes and glial cells at the protein level. (A) Biaxial plots showing the gating strategy to isolate MLCs from other immune and glial cells. P2Y12 expression distinguished a CD45+CD11b+ mononuclear phagocyte subpopulation that does not express glial cell markers S100β and FABP7 and is consistent with the expected MLC phenotype based on sequencing characterization. (B) The proportion of cells expressing P2Y12 within select immune and glial cell subpopulations. P2Y12 is

highly specific for MLCs, which constitute 50% of all CD45+CD11b+ mononuclear phagocytes (MNPs) in this sample. P2Y12 was expressed on approximately 15% of satellite glial cells (SGCs) and was negligibly expressed on other cell types. (C) The frequency of immune and glial cell types as a proportion of all live cells.

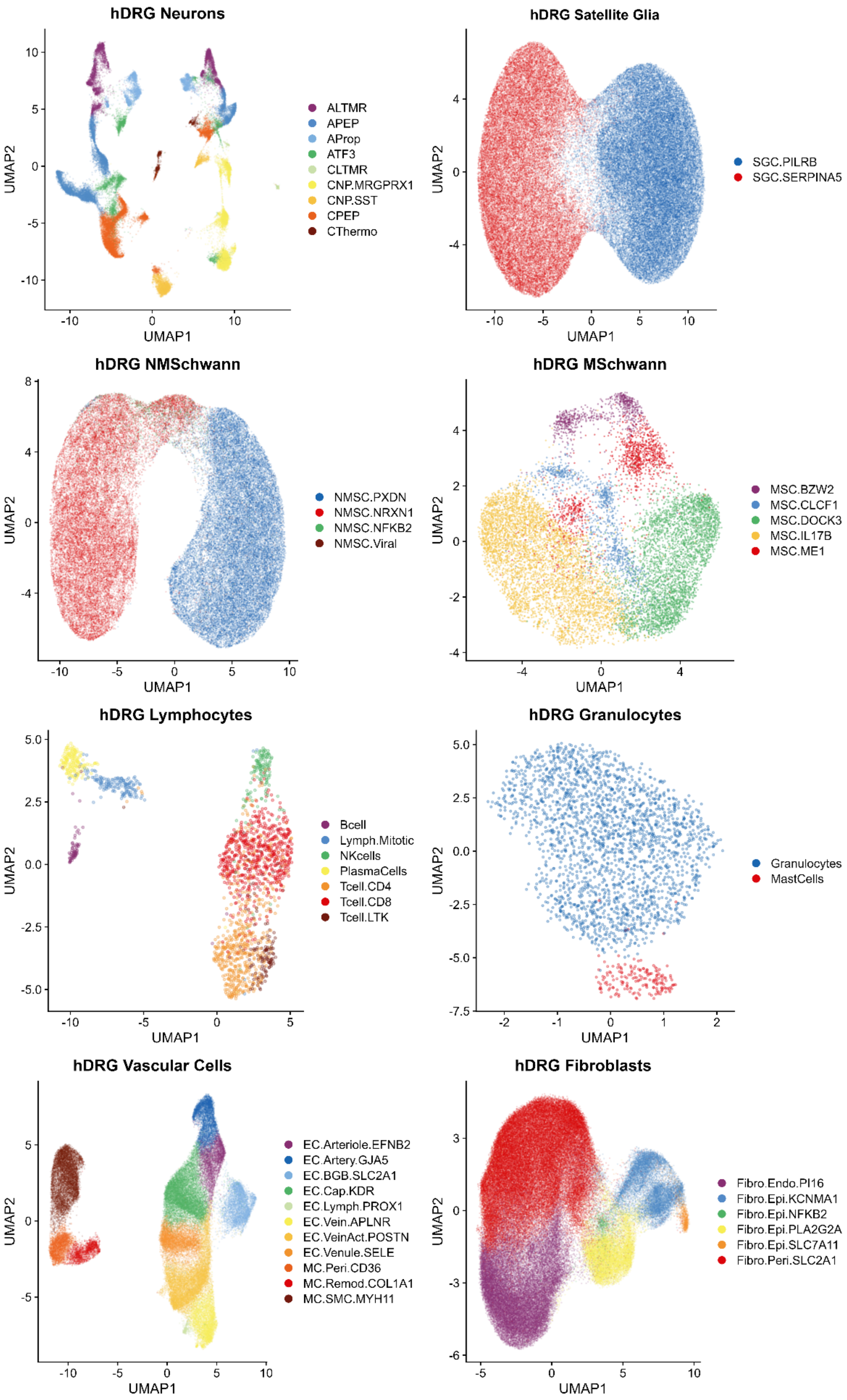


**Figure S3. Other hDRG cell types**

High level cell types from Figure 1A were subclustered to identify a variety of cell types in the hDRG. Only adipocytes were not subclustered. Some clusters (Lymph.Mitotic, NMSC.NFKB2, NMSC.Viral, and Fibro.Epi.NFKB2) may represent cell states rather than distinct cell types.

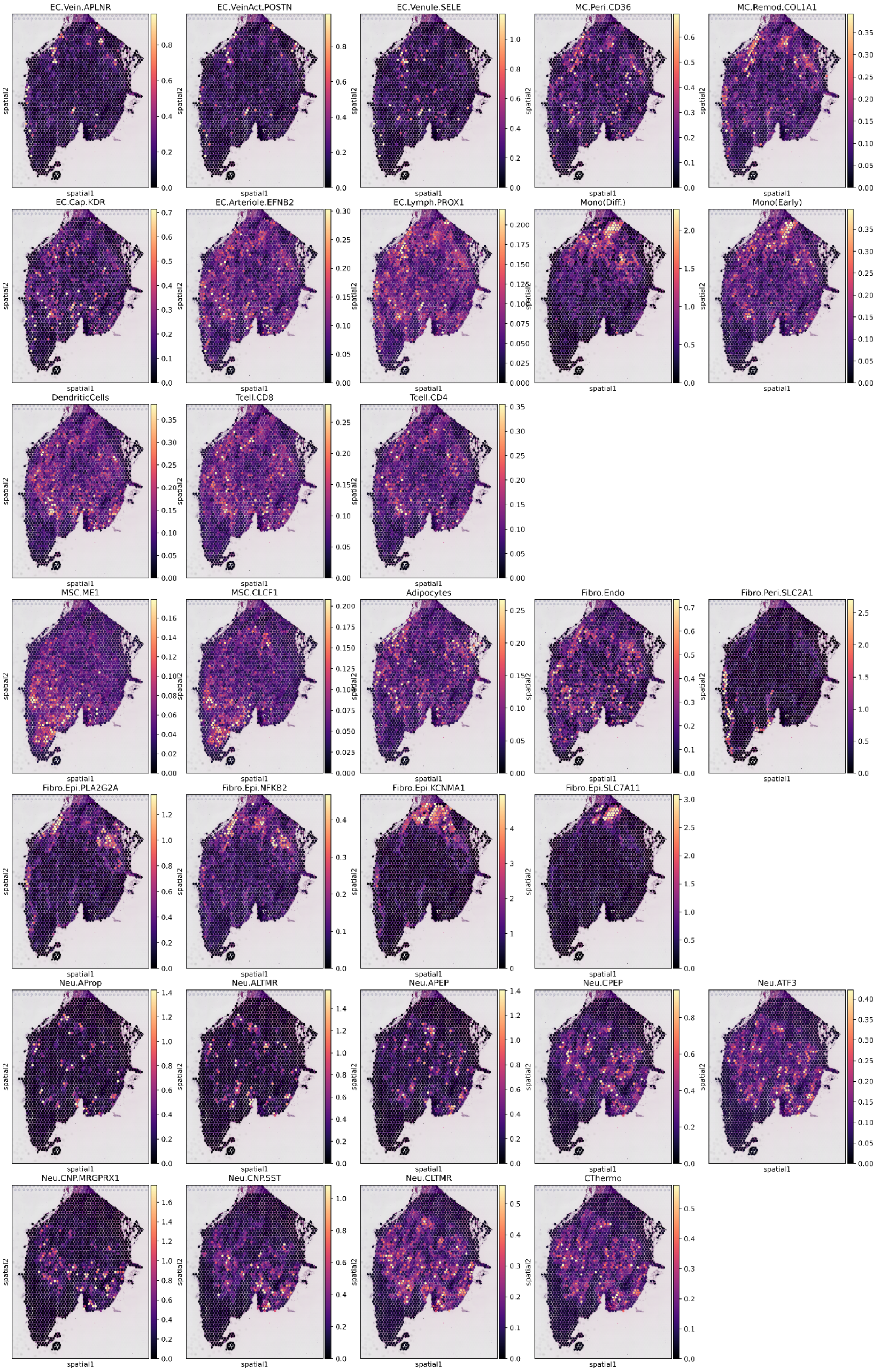

**Figure S4. Predicted localization of hDRG cell types**
Spatial deconvolution exhibited localization patterns among hDRG cell types. Abundance scores per spot for each cell type are shown, with color scaling adjusted per cell type.

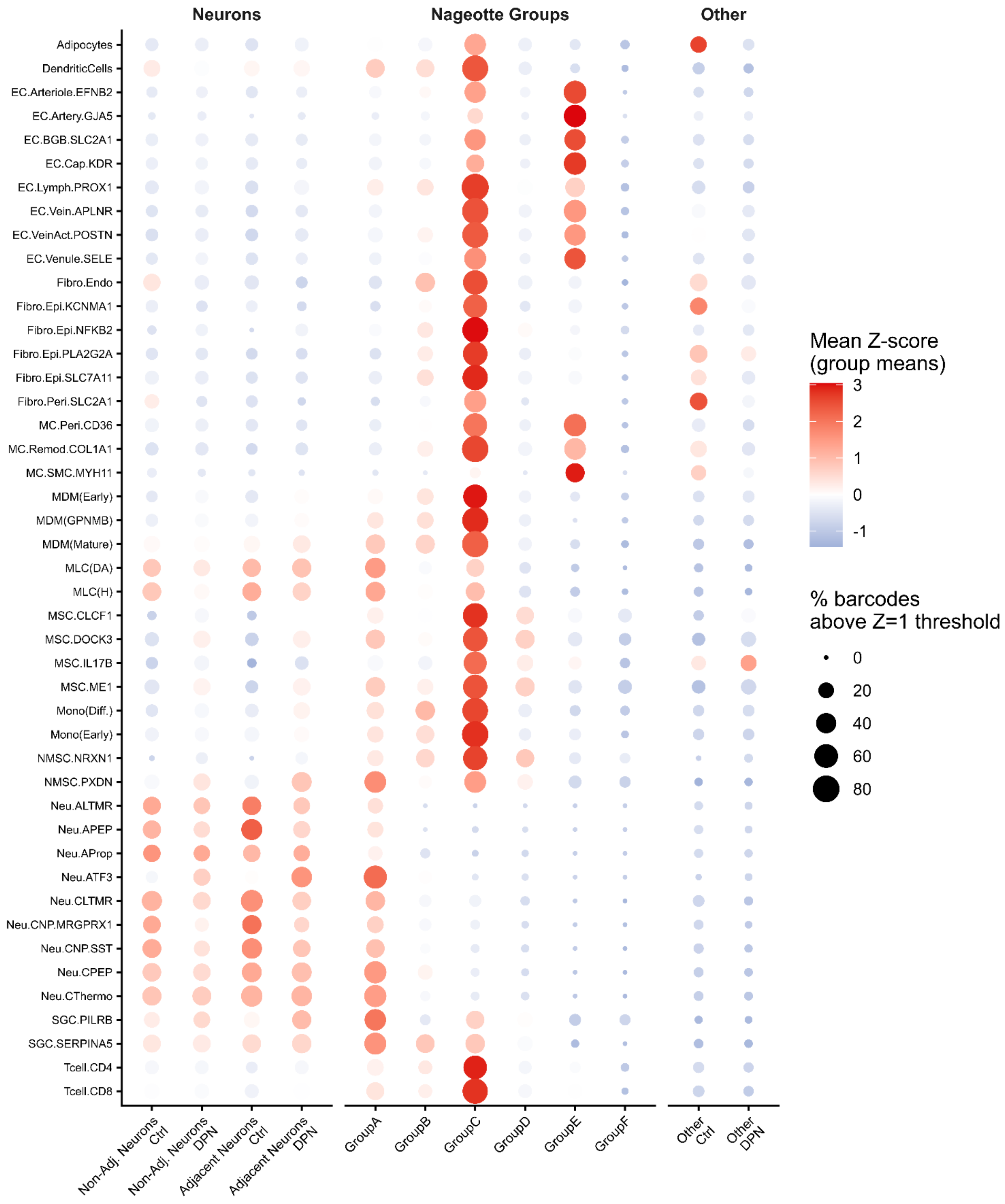


**Figure S5. Enrichment of all non-neuronal hDRG cell types across neuronal and Nageotte nodule spatial barcodes**
Average abundance score for each cell type per barcode type was calculated. For each cell type, these values were scaled by z-score across all barcode types (color) and the percent of barcodes with abundance score greater than z = 1 was tabulated (dot size). Satellite cells including SGCs, MLCs, and PXDN NMSCs were most abundant in neuronal and Group A and C nodule barcodes. Other immune cells and glia were most abundant in Group C nodule barcodes. Fibroblasts were most abundant in Group C nodule and other barcodes. Vascular cells were most abundant in Group C and E nodule barcodes.

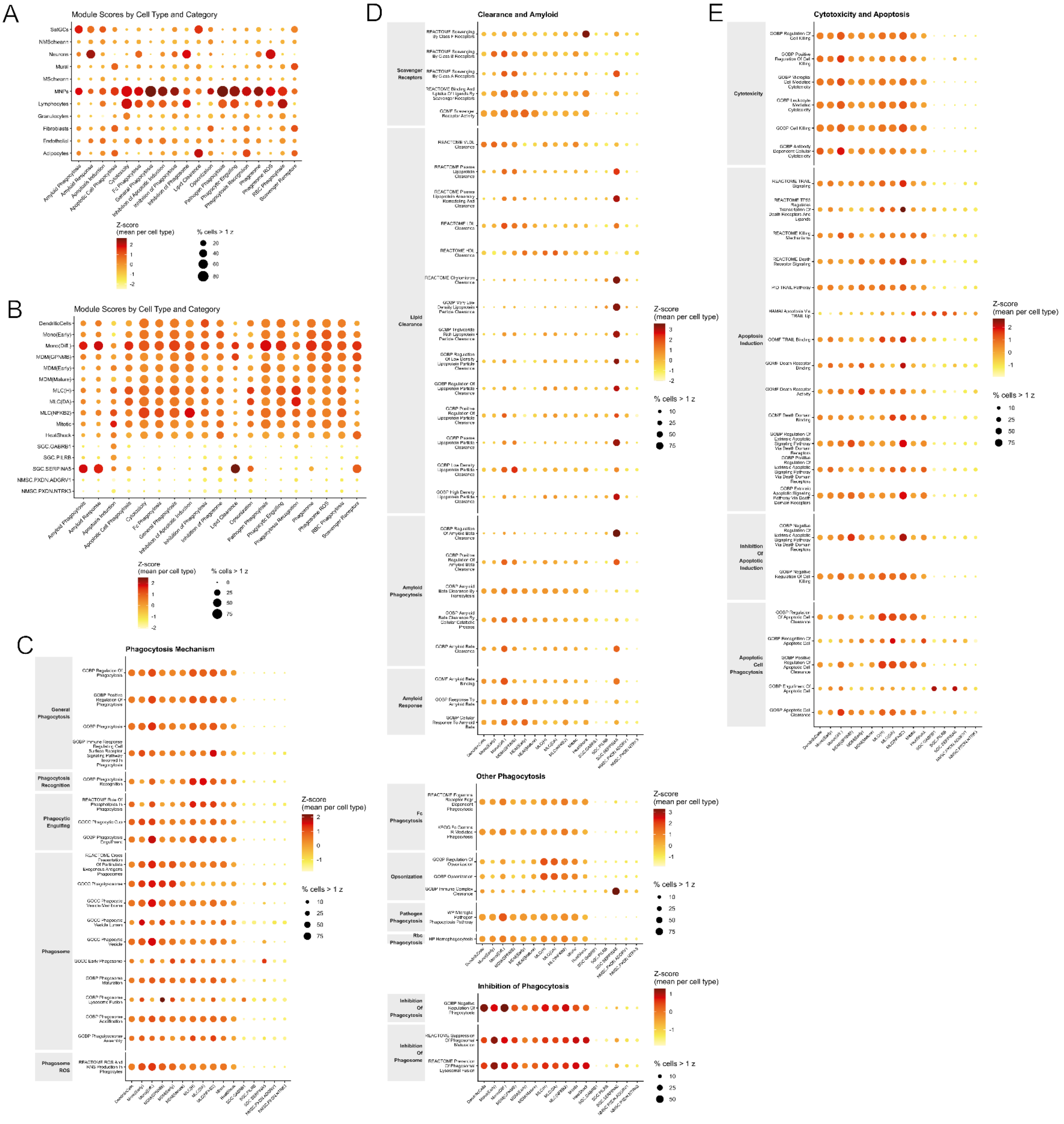


**Figure S6. Enrichment of phagocytosis and apoptosis gene sets in hDRG cells**

A) Enrichment of each of the 15 modules in Figure 5 across all high-level cell types, with color indicating scaled module score (z-score of cell type's average among all cell types) and dot size indicating percent of cells with module score greater than z = 1. B) Enrichment of the 15 modules across MNP and satellite cell types. C-E) Analysis of original phagocytosis and apoptosis gene sets across MNP and satellite cell types.
